# Regional and Systemic Metabolic Remodeling Promotes Longevity by Bioengineered Yeast-Derived Lipids

**DOI:** 10.64898/2026.04.10.717033

**Authors:** Yajuan Li, Zhiliang Bai, Yuhan Li, Fangyuan Gao, Shuo Qin, Jian Ran, Jorge Villazon, Anna Wang, Hongje Jang, Zhi Li, Shriya Sankaran, Yuting Liu, Dorota Skowronska-Krawczyk, Nan Hao, Rong Fan, Lingyan Shi

## Abstract

Aging is marked by a progressive breakdown of intestinal integrity and metabolic homeostasis, which together drives systemic decline in physiology and reduces lifespan. Here, we found that dietary lipids extracted from a genetically engineered long-lived yeast strain robustly extend lifespan in Drosophila and further uncovered the mechanisms using deuterium oxide–probed stimulated Raman scattering microscopy to image metabolic dynamics and single nucleus RNA sequencing (scRNA-seq) to unveil the underlying pathways. These yeast lipids are enriched in shorter, more saturated fatty acids and phospholipids as revealed by Raman spectroscopy and lipidomics, contributing to increased membrane order and reduced lipid storage. Functionally, targeted dietary supplementation with these lipid components synergistically prolongs fly lifespan. We show that these lipids reverse age-related declines in gut lipid droplet abundance, enhance membrane lipid incorporation, and increase de novo lipid synthesis, thereby improving epithelial structural integrity and barrier function. snRNA-seq identifies transcriptional remodeling in metabolically active enterocytes, including upregulation of autophagy and protein turnover genes, alongside reduction of unsaturated fatty acid biosynthesis. In the brain, dietary lipids orchestrate a dual metabolic strategy—promoting energy conservation and enhanced signaling across most neuronal and glial populations, while selectively boosting mitochondrial function in memory-critical Kenyon cells. All these leads to the enhancement of gut-to-glia communication, particularly through EGFR and FGFR pathways. Finally, analysis of our data with the Fly Metabolic Analysis Pipeline (FLY-MAP) reveals that yeast lipids restructure gut metabolic modules to coordinate energy production, redox balance, and nutrient flexibility. Our study uncovers a cross-kingdom mechanism of metabolic longevity regulation, paving the way for leveraging yeast-derived nutritional components to support tissue homeostasis and promote healthy aging.

## Main

Aging is marked by the gradual accumulation of molecular and cellular damage, resulting in widespread physiological decline across the organism^1,2^. This complex process is shaped by both genetic and environmental factors. As a key interface with the environment, the intestine must rapidly adapt to fluctuating conditions such as feeding, starvation, and exposure to toxins or pathogens^3–5^. Maintaining intestinal structural integrity is essential for organismal health under these stresses. Recent studies highlight that age-related metabolic remodeling—alongside declines in both cellular and extracellular intestinal functions and shifts in cytokine expression—is closely linked to the preservation of intestinal integrity during aging^6–9^. Elucidating the molecular mechanisms driving these metabolic shifts is critical for developing strategies to prevent age-associated diseases and promote healthy aging.

Despite its anatomical simplicity, the *Drosophila* midgut serves as a powerful intestinal model, recapitulating many of the cell types and functions found in the mammalian stomach and small intestine^3,10^. Its short lifespan and ease of maintenance enable straightforward longitudinal studies of gut aging, making *Drosophila* an ideal system for exploring the relationship between intestinal physiology and aging^3^. Recent research has shown that dietary components and gut microbiota can influence mortality by modulating gut metabolism and morphology, ultimately affecting lifespan^11–15^. Among various metabolic pathways, lipid metabolism plays a central role in maintaining systemic homeostasis. Aging is associated with a decline in lipid droplet (LD) abundance^16,17^, whereas LD accumulation occurs in response to pathogenic challenges in the *Drosophila* gut^18^. Furthermore, lipid-related dysfunctions caused by genetic mutations in the midgut can be mitigated by dietary lipid supplementation^19,20^. However, how to effectively modulate lipid metabolism to promote longevity in *Drosophila* remains poorly understood.

As a primary dietary source for *Drosophila*, the yeast *S. cerevisiae* provides lipids such as triacylglycerols, steryl esters, phospholipids, and free fatty acids^21–23^. Beyond serving as energy stores, these lipids are essential structural components of cellular membranes. We recently engineered a long-lived yeast strain by co-overexpressing Sir2 and Hap4^24^. Sir2, a founding member of the conserved sirtuin family, promotes gene silencing and genomic stability through lysine deacetylation^25^, while Hap4 is a key regulator of heme biogenesis and mitochondrial function via its role in the heme activator protein (HAP) complex^26^. Since both sirtuins and mitochondria are deeply involved in lipid metabolism across eukaryotes^27,28^, we hypothesized that this engineered yeast strain harbors an altered lipid profile. We further postulated that supplementing *Drosophila* with lipids derived from this long-lived yeast would influence gut physiology and potentially extend lifespan.

In this study, by integrating deuterium oxide (D₂O) probed stimulated Raman scattering (DO-SRS) microscopy imaging, single-nucleus RNA sequencing (snRNA-seq), and mass spectrometry-based lipidomics, we systematically evaluated the gut function and lifespan of flies fed with lipids extracted from the engineered yeast strains. The mechanism underlying this pro-longevity effect involves enhanced de novo lipid synthesis, preservation of gut epithelial integrity, transcriptional reprogramming toward autophagy and proteostasis, and lipid remodeling toward shorter and more saturated species. These changes collectively support metabolic flexibility and reinforce gut–brain communication through glial signaling pathways. Our results reveal a cross-kingdom longevity regulation between two evolutionarily distant organisms and set the stage for developing and using yeast-derived nutritional supplements to promote healthy aging in animals.

## Results

### Long-lived yeast lipids promote gut health and longevity in *Drosophila*

We previously demonstrated that co-overexpression of Sir2 and Hap4 (strain 880) markedly extends yeast lifespan, more so than overexpression of Sir2 (strain 897) or Hap4 (strain 868) alone^24^. Building on this, we investigated whether lipids extracted from these long-lived yeast strains could influence lipid metabolism and lifespan in *Drosophila*. We compared the lifespans of *W^1118^* flies fed with lipid extracts from long-lived yeast strains 880, 868, and 897, with those fed extracts from the wild-type (WT) strain 270 as a control, as well as a separate control group fed standard food without lipid supplementation. Flies fed with 880-derived lipids showed significant increases in both median lifespan (defined as the age at which 50% of the population remains alive) and maximum lifespan (top 10% survivors) (Fig. 1a, b). In contrast, 868-fed flies exhibited only a modest lifespan extension, while no significant changes were observed in flies fed lipids from strain 897, wild-type 270, or standard food.

**Fig. 1:**
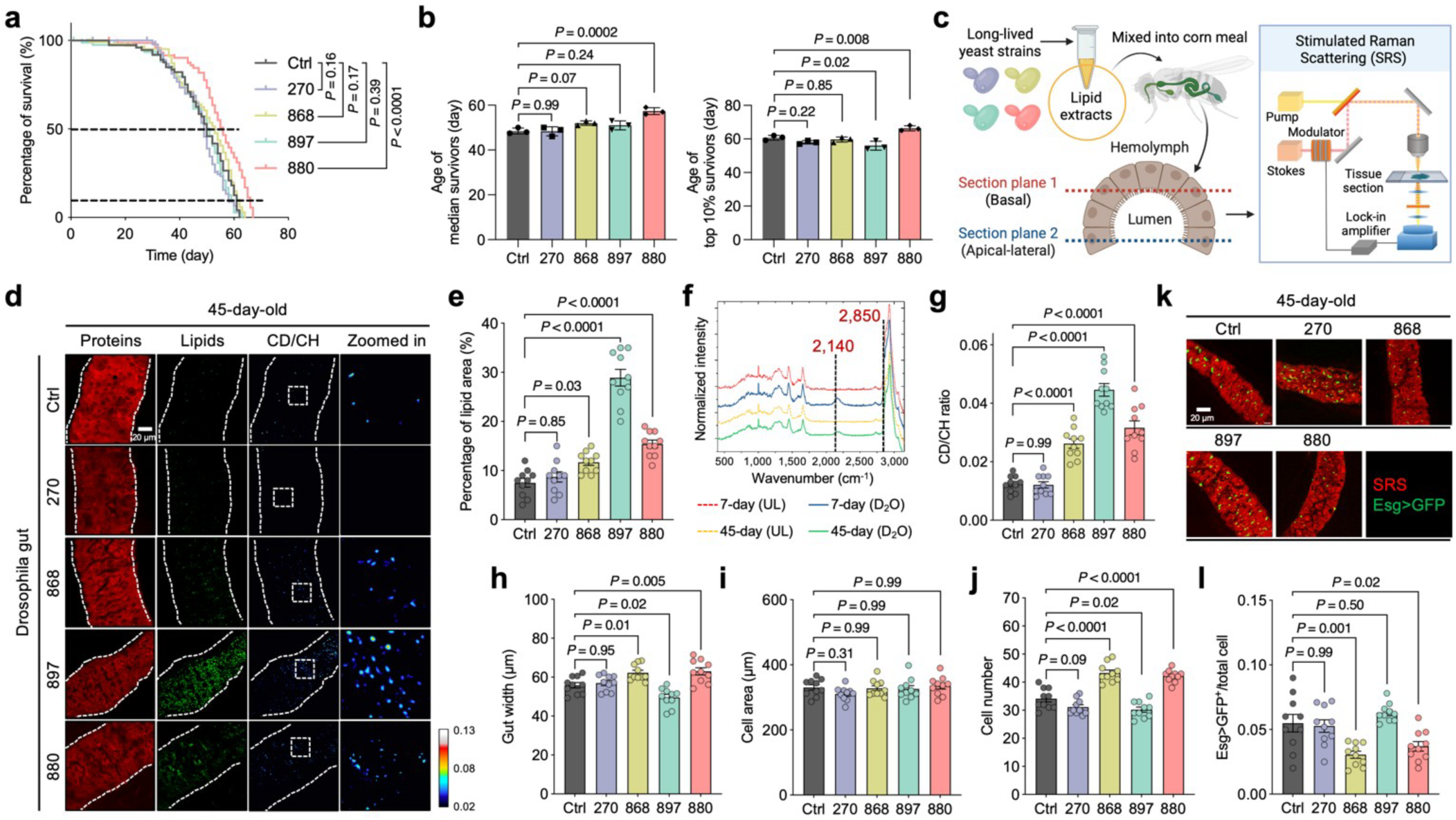
Yeast lipids from long-lived strains promote gut integrity and extend lifespan in *Drosophila*. **a**, Lifespan analysis of flies fed with lipid extracts derived from yeast strains 270, 868, 897, 880, or a blank control. *n* = 120 flies per group. **b**, Comparison of fly age at median survival (left) and top 10% survival (right) across different yeast extract groups. **c**, Schematic illustration of SRS imaging workflow on gut epithelial tissue sections from flies fed with yeast-derived lipid extracts. Created with BioRender.com. **d**, DO-SRS imaging of metabolic profiles in the guts of 45-day-old flies fed with different lipid extracts. CD/CH (2,140/2,850) indicates newly synthesized lipids. **e**, Quantification of total lipid area from panel **d** and comparison across yeast strain groups. **f**, Raman spectra revealing a distinct CD peak in the cell-silent region of the gut from flies fed with D₂O-labeled food versus unlabeled (UL) food. Each spectrum represents an average from *n* = 10 flies. **g**, Quantification of CD/CH (2,140/2,850) ratio in panel **d**, compared across yeast strain groups. **h**–**j**, Measurements of gut width (**h**), average cell area (**i**), and total cell number (**j**) in 45-day-old flies fed with different lipid extracts. **k**, Representative images of Esg>GFP-labeled intestinal stem cells from 45-day-old flies across different lipid extract groups. **l**, Quantification of Esg>GFP⁺ cells as a proportion of total gut cells in panel **k**. Scatter plot shows mean ± s.e.m. from *n* = 3 independent experiments (**b**) and *n* = 10 flies (**e**, **g**–**j**, and **l**). Significance levels were calculated with Log-rank Mantel-Cox test (**a**), or one-way ANOVA with Tukey’s multiple comparisons test (**b**, **e**, **g**–**j**, and **l**).

To visualize lipid distribution in the Drosophila gut, we established a stimulated Raman scattering (SRS) microscopy system to image in situ lipid metabolism with high spatial resolution^29,30^ (Fig. 1c). Previous studies have shown that LDs are primarily localized to enterocytes in the midgut region^3,12^. We therefore focused our imaging on both the basal and apical-lateral sides of gut epithelial cells enriched in enterocytes. Under normal conditions, the lipid signal detected by SRS at the 2,850 cm^−1^ vibrational frequency was predominantly localized to LDs within gut epithelial cells^31^. The specificity of this LD signal was confirmed using conventional lipid staining with BODIPY 493/503 (Extended Data Fig. 1a). Consistent with prior findings^12,32^, SRS imaging revealed that in young (7-day-old) flies, LDs were evenly distributed along the basal and lateral surfaces of the polygonal enterocytes in the midgut (Extended Data Fig. 1b). In contrast, in aged (45-day-old) flies, LD abundance was significantly reduced (Extended Data Fig. 1b, c). These results highlight age-associated changes in lipid distribution, affecting both LDs and membrane-like structures within the gut epithelium.

We next evaluated the effects of dietary lipid extracts on gut lipid metabolism. Compared to flies fed lipids from the wild-type strain 270 or standard food (control), supplementation with lipids from strains 880, 868, or 897 significantly increased LD abundance in the guts of both middle-aged (25-day-old) (Extended Data Fig. 2a, b) and old-aged (45-day-old) flies (Fig. 1d, e). To determine whether the lipids stored in LDs were newly synthesized or translocated from other cellular compartments or tissues, we fed flies D₂O-labeled standard food with or without yeast lipid supplementation. The deuterium (D) from D^2^O is enzymatically incorporated into lipids, replacing hydrogen (H) in carbon-hydrogen (CH) bonds to form carbon-deuterium (CD) bonds, which generate a Raman peak at 2,140 cm^−1^ in the cell-silent region of the spectrum—a known marker of de novo lipid synthesis bonds^29,30^ (Fig. 1f). This CD peak intensity was substantially reduced in the gut of 45-day-old flies compared to 7-day-old young flies. Notably, flies fed lipids from long-lived yeast strains (880, 868, 897) exhibited significantly higher CD/CH (2,140/2,850) ratios in gut tissues on both day 25 (Extended Data Fig. 2c) and day 45 (Fig. 1g), indicating enhanced de novo lipid synthesis. In contrast, lipid synthesis in the fat body, the primary lipid storage organ in *Drosophila*, was unaffected by dietary lipid supplementation (Extended Data Fig. 2d, e).

We also examined the impact of different lipid extracts on gut morphology in aged flies. Gut width increased in flies treated with lipids from strains 868 and 880, but decreased in those treated with 897-derived lipids (Fig. 1h). Gut dimensions are influenced by both cell size and cell number^10^. While cell size remained largely unchanged across groups (Fig. 1i), we observed a significant increase in the number of polygonal enterocytes in the 868 and 880-treated groups (Fig. 1j). In the adult *Drosophila* midgut, intestinal stem cell (ISC) gives rise to absorptive enterocytes^3,33^, prompting us to assess ISC activity using esg-Gal4 > UAS-GFP labeling^17^ (Fig. 1k). The relative ISC counts (normalized to total cell number) were significantly reduced in the 868 and 880 groups, while no change was observed in the 897 group (Fig. 1l). Given that aging is associated with ISC over-proliferation and enterocyte apoptosis—processes that contribute to gut dysplasia and loss of integrity^10,34^—our findings suggest that 868 and 880 lipids help preserve gut homeostasis in aged flies, as evidenced by increased enterocytes and decreased ISCs at day 45. Together, these results demonstrate that lipid extracts from long-lived yeast strains modulate gut lipid metabolism and morphology, with strain 880 exerting the most pronounced effect on promoting longevity.

### Elevated lipid synthesis in long-lived yeast supports gut epithelial membrane integrity in flies

We next characterized the metabolic activity of the long-lived yeast cells using spontaneous Raman microscopy, which provides vibrational signatures specific to molecular structures^35^. Yeasts were cultured in D_2_O-labeled media, and Raman spectra were collected (Fig. 2a). The whole spectra were dominated by peaks at 2,935 cm^−1^, corresponding to CH₃ vibrational modes from proteins, and 2,150 cm^−1^, representing newly synthesized macromolecules in the cell-silent region (Extended Data Fig. 3a). These features indicate active protein content and ongoing macromolecular synthesis during D₂O incubation. We quantified the lipid-to-protein ratio (CH₂/CH₃, 2,850/2,935) and observed a significant increase in the long-lived strains, with the strongest effects seen in strains 868 and 880 (Fig. 2b, c). To further examine lipid biosynthesis, we extracted Raman spectra of deuterium-labeled lipids (D-lipids), which exhibited a distinct peak at 2,176 cm^−1^—separate from the general CD signal at 2,150 cm^−1^ observed in whole-cell spectra (Fig. 2d and Extended Data Fig. 3b). The D-lipid to total lipid ratio (CD/CH, 2,176/2,850) was significantly elevated in all three long-lived strains, indicating enhanced lipid synthesis (Fig. 2e).

**Fig. 2:**
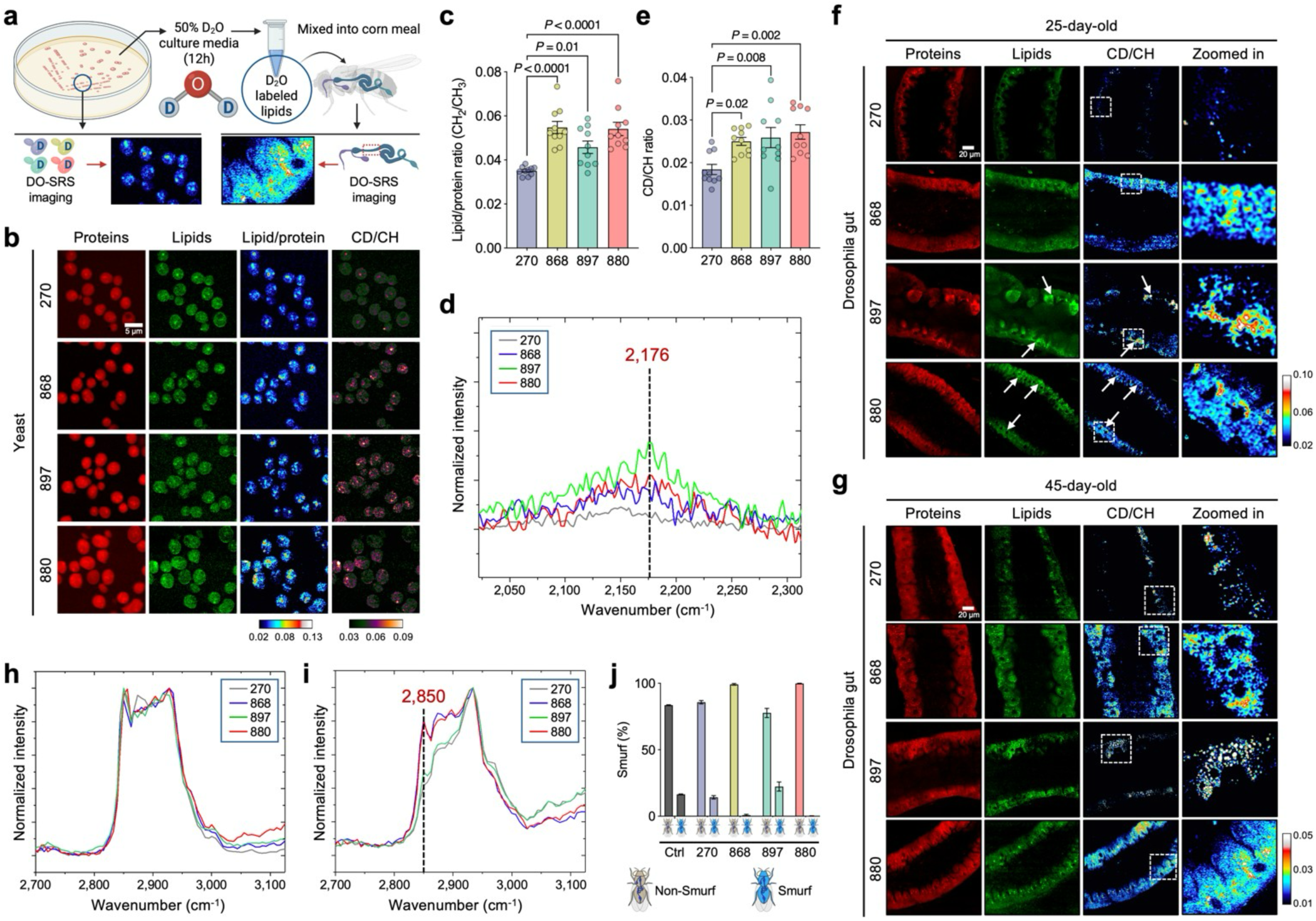
Elevated lipid synthesis in long-lived yeast enhances gut barrier function in flies via membrane incorporation. **a**, Schematic illustration of DO-SRS imaging workflow applied to long-lived yeast cells cultured in D₂O-labeled media and gut epithelial tissue sections from flies fed with D₂O-labeled yeast-derived lipid extracts. Created with BioRender.com. **b**, DO-SRS imaging of metabolic profiles in yeast cells. CD/CH (2,176/2,850) indicates D-lipids. **c**, Quantification of lipid/protein ratio from panel **b** and comparison across yeast strain groups. **d**, Raman spectra of the CD region showing a distinct D-lipid peak at 2,176 cm⁻¹. Each spectrum represents an average from *n* = 10 flies. **e**, Quantification of CD/CH (2,176/2,850) ratio in panel **b**, compared across yeast strain groups. **f, g**, DO-SRS imaging of gut metabolic profiles in 25-day-old (**f**) and 45-day-old (**g**) flies fed with different D₂O-labeled yeastderived lipid extracts. **h**, **i**, Hyperspectral Raman spectra of lipid droplets (**h**) and membranes (**i**) from fly guts shown in panel **f**. Each spectrum is averaged from *n* = 10 region of interests. **j**, Smurf assay assessing gut barrier integrity in flies fed with lipid extracts from different yeast strains. *n* = 80 flies from 4 independent experiments. Scatter plot shows mean ± s.e.m. from *n* = 10 flies (**c**, **e**). Significance levels were calculated with one-way ANOVA with Tukey’s multiple comparisons test (**c**, **e**).

Barrier function is closely linked to the composition and abundance of membrane lipids^36^. To assess whether exogenous lipids from yeast are incorporated into the gut epithelium, we fed flies with lipid extracts from D₂O-labeled yeast strains and performed DO-SRS imaging at 2,176 cm^−1^ (Fig. 2a), a peak corresponding to D-lipids. On day 25, flies fed with 880- and 868-derived D-lipids displayed a well-organized, single-layered columnar epithelium with aligned nuclei and uniform cytoplasmic distribution of D-lipids (Fig. 2f). In contrast, flies fed with D-lipids from the WT strain (270) exhibited disorganized, tumor-like epithelial structures and notably lower CD signal compared to other groups. Interestingly, in 897-fed flies, D-lipids accumulated in LD-like structures and were associated with irregular epithelial morphology, suggesting altered lipid homeostasis distinct from that seen in the 868 and 880 groups. By day 45, the membrane lipid architecture in 868- and 880-fed flies remained largely intact (Fig. 2g), resembling that of 7-day young flies (Extended Data Fig. 1b), highlighting the sustained protective effects of these lipid extracts on gut structural integrity.

Hyperspectral SRS imaging revealed minimal differences in the spectral shape of LDs across the different lipid groups (Fig. 2h), suggesting that the core composition of storage lipids remained consistent. However, the lipid signal, reflected by the Raman peak at 2,850 cm^−1^, was notably elevated in membrane structures of gut epithelial cells in flies fed 880 and 868 lipids (Fig. 2i), indicating increased incorporation of these lipids into cytoplasmic and plasma membranes. These findings suggest that lipids from 880 and 868 yeast strains may enhance gut epithelial integrity by contributing to membrane biosynthesis while supporting LD metabolic homeostasis. To further assess gut barrier function, we performed a Smurf assay in aged (45-day-old) flies, in which flies co-ingest a blue dye that normally does not cross the intestinal barrier^16^. An average of 16%, 14%, 1%, and 22% of flies in the control, 270-fed, 868-fed, and 897-fed groups exhibited the Smurf phenotype (Fig. 2j), indicating dye leaking to the body cavities. In contrast, no Smurf phenotype was observed in flies fed with 880 lipids, suggesting preserved barrier integrity in these groups. Taken together, these results demonstrate that long-lived yeast strains exhibit elevated lipid synthesis and that these lipids can be effectively incorporated into gut epithelial membranes to enhance gut barrier function, an effect that may contribute to lifespan extension in *Drosophila*.

### snRNA-seq reveals alterations in autophagy, protein turnover, and lipid biosynthesis in 880-fed fly guts

To investigate how lipid products from engineered yeasts influence gut physiology and lifespan, we performed snRNA-seq on gut tissues from 45-day-old flies fed with lipid extracts from strains 880, 868, or a control diet without lipid supplementation, strictly following the Fly Cell Atlas protocol^37,38^ (Fig. 3a). Paired brain tissues were also collected, given the critical role of the gut-brain axis in Drosophila aging^39,40^. Despite higher sequencing saturation in brain samples, gut tissues showed higher gene counts (Extended Data Fig. 4a), consistent with observations from the Fly Cell Atlas. After removing low-quality nuclei and potential doublets, we retained 32,804 gut and 38,277 brain high-quality nuclei for analysis. Unsupervised clustering and Uniform Manifold Approximation and Projection (UMAP) embedding of the integrated dataset identified 20 distinct subpopulations (Extended Data Fig. 4b), each defined by unique transcriptomic profiles and marker gene expression (Extended Data Fig. 4c). Clusters were primarily segregated by tissue type, and samples from different conditions exhibited consistent distributions, indicating minimal batch effects (Extended Data Fig. 4d). Moreover, canonical marker genes for gut and brain were clearly separated on the integrated UMAP (Extended Data Fig. 4e), supporting the accuracy and robustness of our dataset.

**Fig. 3:**
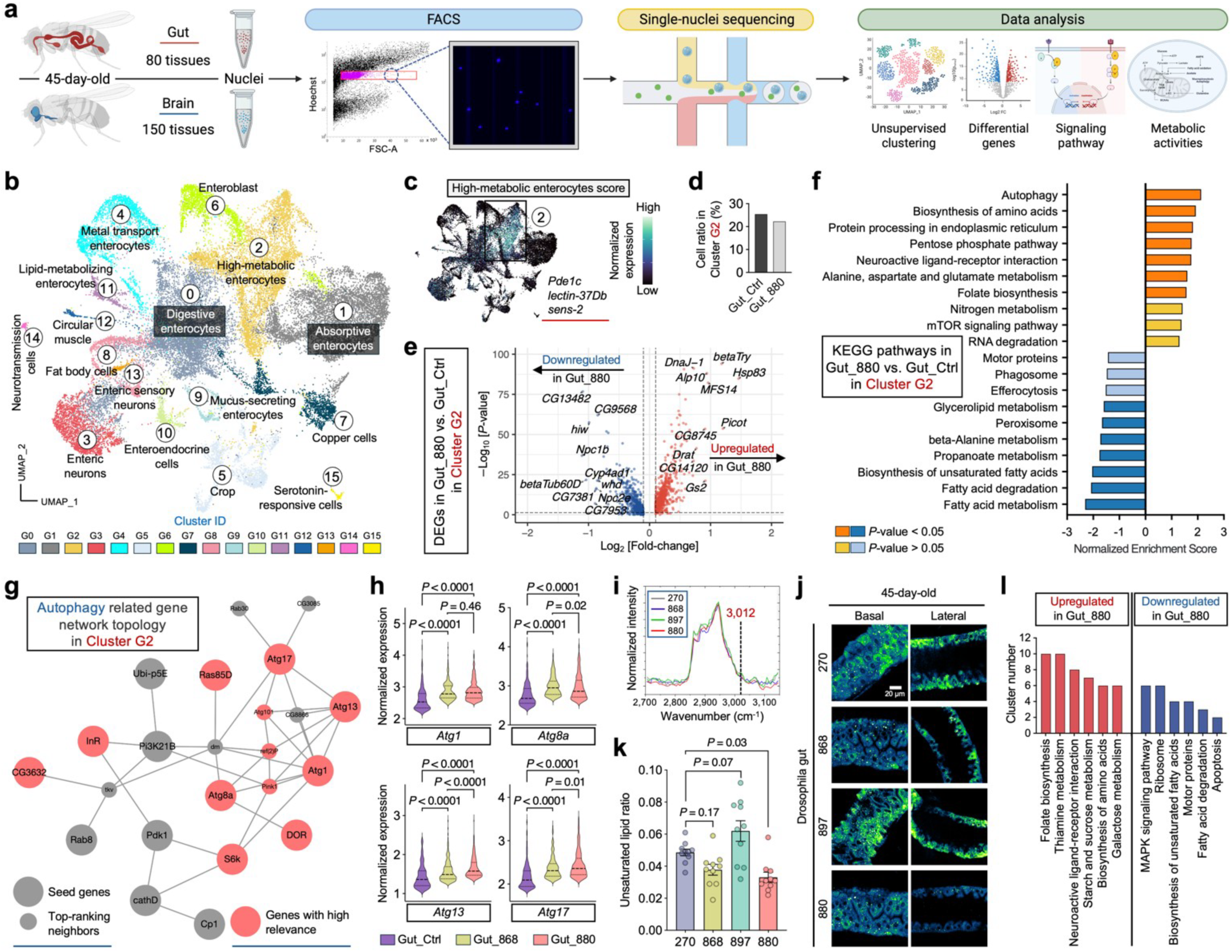
snRNA-seq reveals cell type-specific metabolic reprogramming in 880-fed fly guts. **a**, Schematic of the snRNA-seq experimental design. Created with BioRender.com. **b**, Unsupervised clustering of gut single nuclei from flies fed with lipid extracts from strains 880, 868, or a control (Ctrl) diet without lipid supplementation, with cell type annotations indicated on the UMAP. **c**, Expression distribution of high-metabolic enterocytes score on the UMAP. Genes defining this module score are listed below. **d**, Comparison of cell ratios in cluster G2 between 880-fed flies and control group. **e**, DEGs in cluster G2 nuclei of 880-fed flies compared to control flies. **f**, Corresponding KEGG signaling pathways regulated by the DEGs identified in panel **e**. **g**, Network topology analysis of autophagy-related genes in cluster G2. **h**, Single-nucleus level expression of key genes identified in panel **g** and comparison across dietary groups. **i**, Raman spectra highlighting the peak at 3,012 cm⁻¹, characteristic of unsaturated lipids. Each spectrum represents an average from *n* = 10 flies. **j**, Ratiometric images of the 3,012/2,850 signal on both basal and lateral sides of gut epithelial cells from 45-day-old flies fed with different lipid extracts. **k**, Quantification of the unsaturated-to-total lipid ratio (3,012/2,850) from panel **j** and comparison across yeast strain groups. **l**, Top-ranked signaling pathways significantly (*P*-value < 0.05) upregulated or downregulated in 880-fed flies across most identified clusters. Scatter plot shows mean ± s.e.m. from *n* = 10 flies (**k**). Significance levels were calculated with two-tailed Mann-Whitney test (**h**), or one-way ANOVA with Tukey’s multiple comparisons test (**k**).

To dissect gut cellular heterogeneity, we next performed clustering analysis on gut nuclei, revealing 16 subpopulations, including major cell types such as enterocytes, enteroblasts, and enteroendocrine cells (Fig. 3b and Extended Data Fig. 5a). Notably, cluster G2 enterocytes exhibited a high metabolic state, characterized by elevated expression of *Pde1c* (Fig. 3c), a phosphodiesterase gene involved in cAMP and cGMP regulation^41^, highlighting this cluster for further investigation. Although G2 cell proportions were similar between 880 and the control group (Fig. 3d), differentially expressed genes (DEGs) analysis revealed striking differences (Fig. 3e). In 880-fed flies, numerous genes were significantly up- or downregulated relative to controls. To understand the signaling pathways regulated by these DEGs, we performed Kyoto Encyclopedia of Genes and Genomes (KEGG) analysis and found that autophagy was the most upregulated pathway in the 880 group (Fig. 3f), a process previously shown to contribute to *Drosophila* longevity^42^. Network topology analysis^43^ of autophagy-related genes highlighted *Atg1*, *Atg8a*, *Atg13*, and *Atg17*—all key regulators of autophagy initiation^44^—as central nodes (Fig. 3g), with significantly elevated expression in both 880- and 868-fed flies (Fig. 3h). In addition to autophagy, pathways involved in biosynthesis of amino acids, protein processing in the endoplasmic reticulum, alanine, aspartate and glutamate metabolism, and folate biosynthesis were also significantly upregulated (Fig. 3f), suggesting enhanced protein turnover. Given that protein aggregation is a hallmark of aging and age-related diseases^45^, these findings suggest that 880-derived lipids may mitigate age-associated decline by promoting proteostasis in aged gut cells.

Conversely, several lipid metabolism related pathways were significantly downregulated in cluster G2 cells from 880-fed flies, particularly the biosynthesis of unsaturated fatty acids (Fig. 3f). Key genes involved in this pathway, including *Baldspot*, *Acox57D-d*, and *CG17544*, showed significantly lower expression in both 880 and 868 groups (Extended Data Fig. 5b, c). To validate these findings, we analyzed SRS signals at 3,012 cm^−1^ (Fig. 3i), a Raman peak specific to unsaturated lipids^46^, in 45-day-old gut tissues. Consistent with transcriptomic results, the intensity of the unsaturated lipid signal was reduced on both the basal and lateral sides of epithelial cells in the 880 group (Fig. 3j). Quantitatively, the 3,012/2,850 intensity ratio (unsaturated/total lipids) was significantly lower in 880-fed flies (Fig. 3k). This trend was also observed in 25-day-old gut cells based on SRS imaging (Extended Data Fig. 5d). Since membrane unsaturation affects susceptibility to lipid peroxidation and is linked to aging rates^47^, these data suggest that while 880-fed aged flies maintained high levels of total LDs, their lower levels of unsaturated fatty acids may have reduced vulnerability to lipid peroxidation-related damage.

DEGs and pathway analysis across all clusters consistently revealed downregulation of unsaturated fatty acid biosynthesis in 880-fed flies, particularly in cluster G1 (absorptive enterocytes, ∼20% of total cells), G14 (neurotransmission cells), and G15 (serotonin-responsive cells) (Fig. 3l and Supplementary Table 1). In parallel, biosynthesis of amino acids was significantly upregulated across five additional clusters, including G1, G6 (enteroblasts), G10 (enteroendocrine cells), G12 (circular muscle cells), and G14, further supporting global enhancement of protein turnover in aged guts.

We also performed a global comparison across all the nuclei between 880 and 868-fed flies and found a significant upregulation of circadian rhythm pathway in the 880 group (Extended Data Fig. 5e), supported by elevated expression of key circadian genes including *Clk*, *Pdp1*, *sgg*, *tim*, and *vri* (Extended Data Fig. 5f, g). As autophagy and circadian regulation are interconnected processes implicated in aging^42,48^, this reduction in circadian gene expression may partly explain why 868-fed flies, despite maintaining lipid homeostasis, did not show the same lifespan extension as 880-fed flies. Collectively, unsupervised clustering of single-nucleus transcriptomics revealed complex cellular heterogeneity in the aging gut. A metabolically active enterocyte subpopulation (cluster G2) displayed enhanced autophagy and protein turnover, alongside reduced biosynthesis of unsaturated lipids in 880-fed flies. These transcriptional changes likely contribute to the improved gut homeostasis and extended lifespan observed in flies supplemented with lipids from yeast strain 880.

### DO-SRS imaging and lipidomics uncover lipid remodeling toward saturation in 880 yeast

Given the distinct lipid saturation levels observed in flies fed with different yeast-derived lipids, we next examined the Raman spectra of D-lipids collected from yeast cells, focusing on the CH stretching region (2,800−3,050 cm^−1^), which reflects chain length and saturation level^49^. Lipids from strains 270, 868, and 897 exhibited similar spectral profiles, which could be roughly divided into four characteristic peaks: ∼2,850 cm^−1^, corresponding to CH_2_ symmetric stretching; ∼2,871 cm^−1^, representing stretching vibrations in CH_2_ groups adjacent to an unsaturated carbon atom [–CH_2_–CH=CH–] present in unsaturated fatty acids^50^; ∼2,927 cm^−1^, corresponding to CH_3_ asymmetric stretching; and ∼2,958 cm^−1^, another CH_3_ asymmetric stretching peak^49^ (Fig. 4a).

**Fig. 4:**
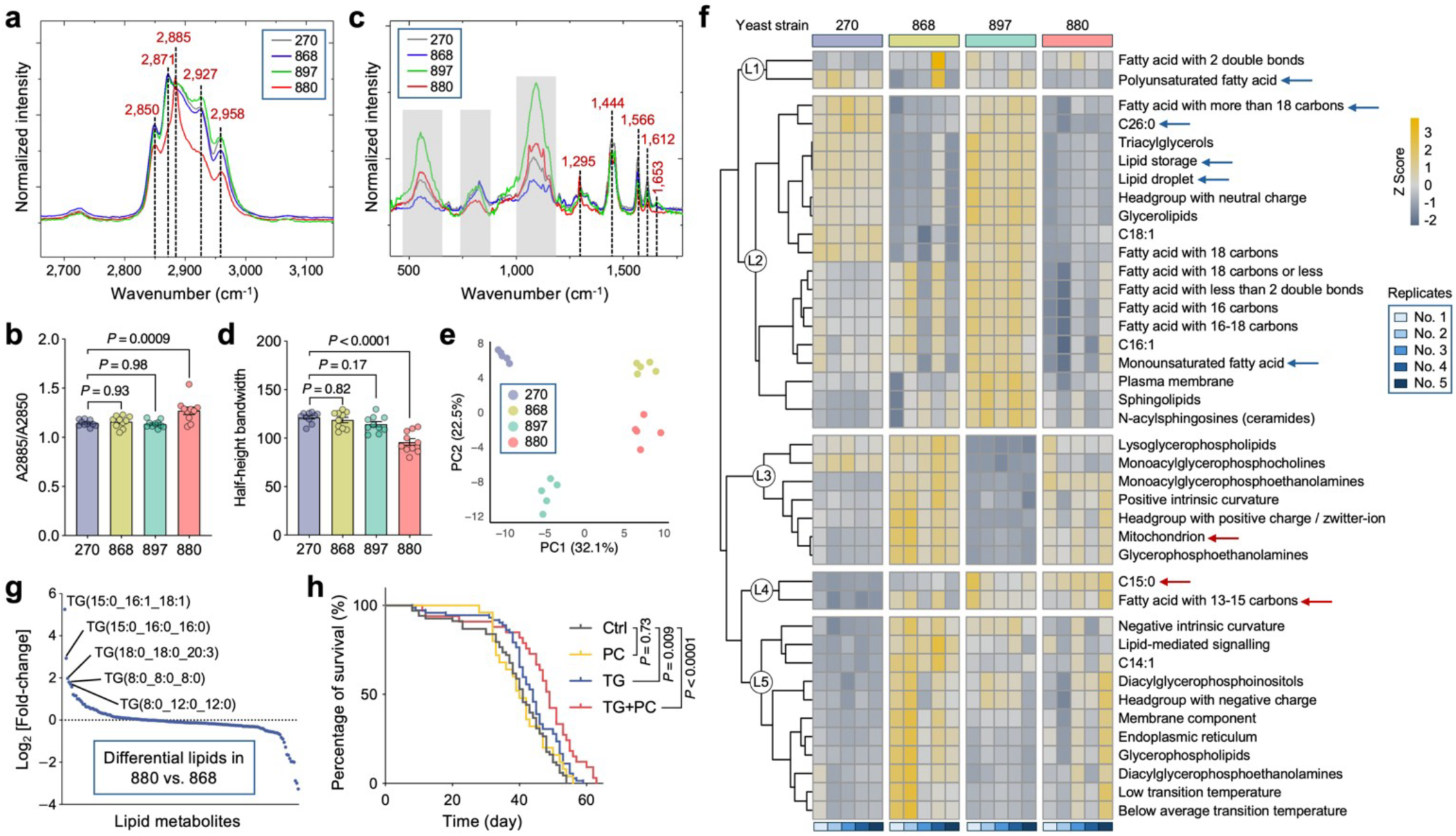
Lipid remodeling in 880 yeast favors saturated, short-chain fatty acids. **a**, Raman spectra of the CH region from D-labeled yeast cell lipids, showing a distinct peak at 2,885 cm⁻¹ in strain 880. **b**, Quantification of the area ratio under the Raman bands at 2,885 cm⁻¹ and 2,850 cm⁻¹ (A2885/A2850) from panel **a** and comparison across yeast strain groups. **c**, Raman spectra of the fingerprint region from D-labeled yeast cell lipids, revealing a narrow peak at 1,295 cm⁻¹ in strain 880. **d**, Quantification of the half-height bandwidth at 1,295 cm⁻¹ from panel **c**, compared across yeast strain groups. **e**, PCA of lipidomic profiles encompassing 196 lipid species from four yeast strains. Each strain includes *n* = 5 replicate measurements. **f**, Lipid Ontology enrichment analysis based on lipidomic expression data. Pathways were clustered into five groups using hierarchical clustering. Z-score > 0 indicates upregulation; Z-score < 0 indicates downregulation. Only significantly enriched pathways (*P*value < 0.05) are shown. **g**, Differential lipid analysis comparing strain 880 to strain 868. Lipid species are ranked by fold-change. **h**, Lifespan analysis of flies fed with medium-chain saturated TG, short- and medium-chain saturated PC, or both. *n* = 120 flies per group. In **a** and **c**, each spectrum represents an average from *n* = 10 flies. Scatter plot shows mean ± s.e.m. from *n* = 10 flies (**b**, **d**). Significance levels were calculated with one-way ANOVA with Tukey’s multiple comparisons test (**b**, **d**).

Notably, 880-derived lipids displayed a distinct spectral feature, with a pronounced peak at 2,885 cm^−1^ (Fig. 4a). This peak is enhanced by Fermi resonance in ordered lipid packing, whereas the 2,850 cm^−1^ band remains unaffected by structural changes in the lipid hydrocarbon chains^51^. Thus, the degree of lipid ordering can be quantified by the ratio of areas under the Raman bands at 2,885 cm^−1^ (A2885) and 2,850 cm^−1^ (A2850)^51^. This ratio was significantly higher in 880-derived lipids (Fig. 4b), indicating increased lipid ordering. Additionally, we analyzed the fingerprint region from 400 to 1,650 cm^−1^, which corresponds to various types of C-C stretching and vibration models. The 880 strain exhibited a distinctly different peak profile compared to other strains (Fig. 4c). In particular, we observed a narrow peak at 1,295 cm^−1^ (associated with the −(CH_2_)n− in-phase twisting mode) in 880 lipids, with a significantly reduced half-height bandwidth (Fig. 4d). This remodeling of the Raman peak shape further supports the presence of highly ordered (CH_2_)n chains^52^, consistent with our findings in the CH stretching region. Altogether, these data strongly suggest that lipids in the 880 yeast strain undergo a phase transition to a more ordered state, in contrast to other strains where lipids likely remain in a more amorphous state.

To further validate the lipid composition, we performed unbiased lipidomics profiling using reversed-phase liquid chromatography–mass spectrometry (RPLC-MS) on lipid extracts from the four yeast strains. This analysis faithfully annotated 196 lipid species spanning 16 major classes (Supplementary Table 2), including acylcarnitines (AcCa), ceramides (Cer), cholesterol esters (ChE), diacylglycerols (DG), lysophosphatidylcholines (LPC), lysophosphatidylethanolamines (LPE), lysophosphatidylserines (LPS), monoglycerides (MG), phosphatidic acids (PA), phosphatidylcholines (PC), phosphatidylethanolamines (PE), phosphatidylglycerols (PG), phosphatidylinositols (PI), phosphatidylserines (PS), triacylglycerols (TG), and wax esters (WE). Principal component analysis (PCA) revealed clear separation among the four strains, suggesting that lipid remodeling occurred in the engineered yeasts relative to the WT strain 270 (Fig. 4e). Strains 880 and 868 clustered closely with high PC1 scores indicating similar lipid profiles, consistent with previous analyses. These two strains showed significantly increased relative abundance of phospholipids, including LPE, PA, and PE (Extended Data Fig. 6a). Phospholipids are fundamental building blocks of all cell membranes, playing a crucial role in both structure and function^53^, which aligns with our previous findings that lipids from strains 880 and 868 enhance gut epithelial integrity in flies (Fig. 2f, g). Moreover, since PS decarboxylation is the primary source of PE in yeast^54^, the observed increase in PE and LPE levels, along with the significant reduction in PS, likely reflects upregulated PE turnover in 880 and 868 yeast cells via the PS–PE–LPE axis.

Unsupervised clustering of the lipidomic profile resolved five groups, reinforcing the similarity between 270 and 897, and between 868 and 880 (Extended Data Fig. 6b). To identify lipid-associated functional pathways within the lipidomes, we performed Lipid Ontology (LION) enrichment analysis^55^, which revealed five clusters primarily separated by chain length, saturation level, molecular shape, and fundamental biological processes (Fig. 4f). Lipids from strain 880 showed notable downregulation of LION pathways including polyunsaturated fatty acid, fatty acid with more than 18 carbons, C26:0, and monounsaturated fatty acid in clusters L1 and L2. We extracted the defining lipid metabolites for each term (Supplementary Table 3), summed their expression levels, and confirmed significantly lower abundance of these pathways in strain 880 (Extended Data Fig. 6c). Furthermore, we observed a significant reduction of lipid storage and LD, along with elevated phospholipid levels, suggesting a shift from neutral lipid storage toward membrane lipid composition. Conversely, C15:0 (pentadecanoic acid), an odd-chain saturated fatty acid, showed significant upregulation in strain 880, along with other medium-chain fatty acid with 13-15 carbons in cluster L4 (Fig. 4f and Extended Data Fig. 6d). Several short-chain saturated lipids were also specifically elevated in the 880 yeast, including LPC(15:0), PE(10:0_14:0), TG(8:0_8:0_8:0), and DG(12:0_14:0) (Extended Data Fig. 6b), further supporting a fatty acid composition shift toward shorter and more saturated species, consistent with our Raman spectroscopy findings.

Considering the high lipidomic similarity between strains 880 and 868, yet only 880-derived lipids significantly extended fly lifespan, we performed differential lipid analysis between these two strains. This revealed that a group of TGs was among the most upregulated lipids in strain 880 compared to 868 (Fig. 4g), although both strains showed overall reduced TG levels relative to the WT 270 strain (Extended Data Fig. 6a). Based on these findings, we hypothesized that this subset of TGs contributes to lifespan regulation. To test this, we fed flies with cornmeal containing three distinct lipid compositions: (1) medium-chain saturated TG (8:0_8:0_8:0) (1,2,3-Trioctanoylglycerol); (2) a mixture of short- and medium-chain saturated phospholipids, including PC(4:0), PC(6:0), and PC(10:0), given that PC is the predominant phospholipid class essential for membrane structure^56^; and (3) a combination of both TG and PC. Feeding flies with TG alone extended lifespan, whereas PC alone did not (Fig. 4h). Remarkably, combined supplementation with both TG and PC, resembling the unique lipidomic signature of strain 880, resulted in significantly greater lifespan extension, suggesting synergistic benefits of these lipid species. Together, these findings demonstrate that the 880 yeast strain undergoes lipid remodeling toward shorter, more saturated fatty acids, enriching membrane-associated phospholipids and enhancing lipid order. This distinct lipid profile is likely a key contributor to enhanced gut epithelial integrity, consistent with our transcriptomic analyses showing improved gut homeostasis in 880-fed flies.

### 880-fed flies exhibit brain metabolic reprogramming and enhanced gut-glial communication

To explore how dietary lipid intervention might impact brain activity in aging flies, we performed clustering analysis of brain nuclei from flies fed with lipid extracts and identified 22 transcriptomic clusters (Fig. 5a and Extended Data Fig. 7a). The distribution patterns on the UMAP confirmed the robustness of the clustering, with clear segregation between neuronal and glial cell clusters, and distinct separation of Kenyon cells (KCs) from other neuronal subtypes. In *Drosophila*, an extensive analysis of single-cell transcriptomes from ageing fly brains has shown that glial cells, but not neurons, exhibit pronounced age-related transcriptional trajectories^57^. Moreover, KCs within the mushroom body, a key brain region for learning and memory, undergo significant age-dependent changes^58^. Therefore, we focused our analyses on two major glial subpopulations, ensheathing glial cells (cluster B3) and perineurial glial cells (cluster B5), along with two KC subtypes, Gamma KCs (cluster B4) and Alpha/beta KCs (cluster B8), for deeper investigation.

**Fig. 5:**
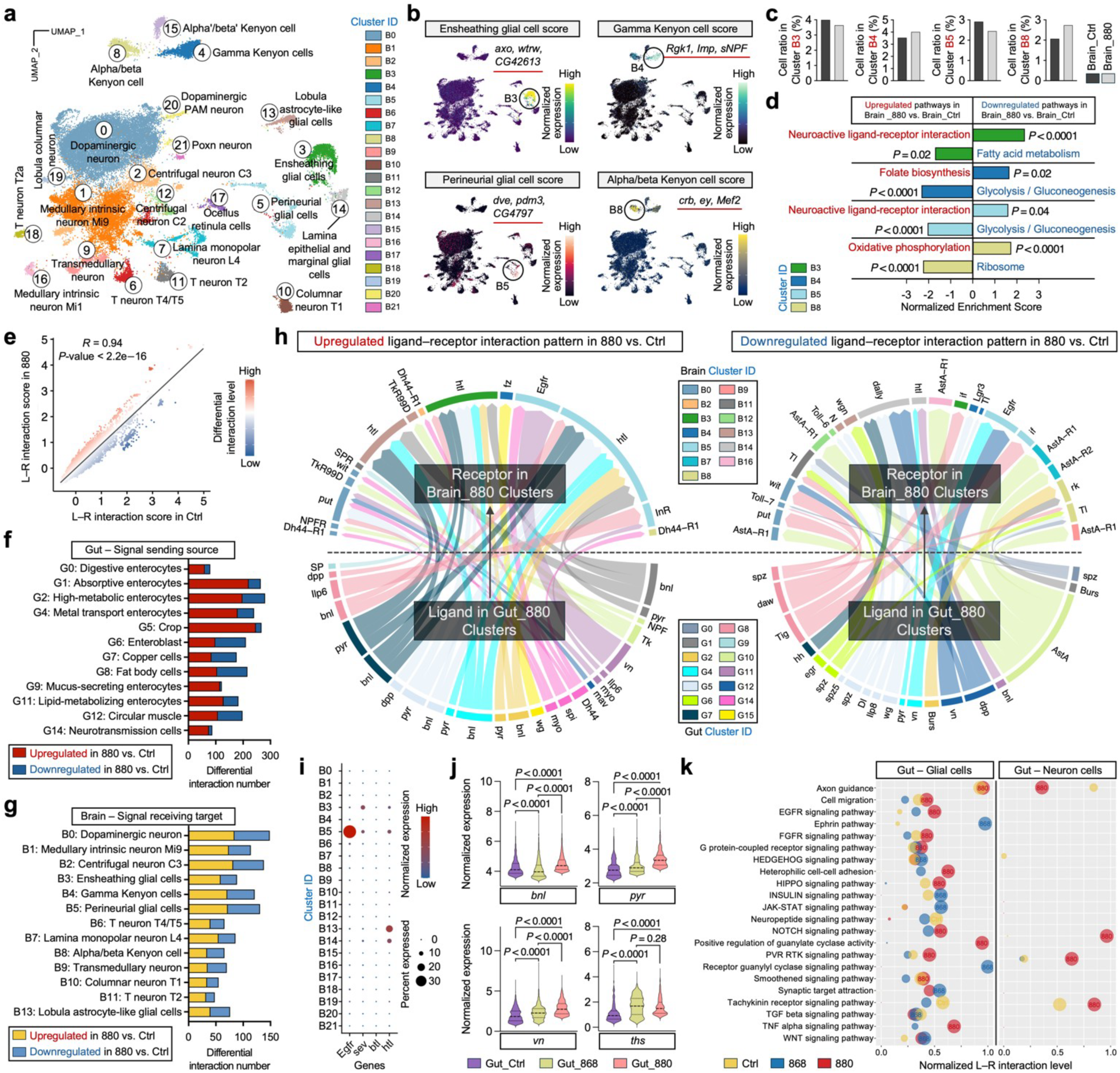
880 lipids modulate brain metabolism and enhance gut-glial communication via EGFR/FGFR signaling. **a**, Unsupervised clustering of brain single nuclei from flies fed with lipid extracts from strains 880, 868, or a control (Ctrl) diet without lipid supplementation, with cell type annotations indicated on the UMAP. **b**, Expression distribution of selected cell type scores on the UMAP. Genes defining each module score are listed below. **c**, Comparison of cell ratios between 880-fed flies and control group across cell type-associated clusters shown in panel **b**. **d**, Top-ranked signaling pathways upregulated or downregulated in 880-fed flies compared to control group across different clusters. **e**, Correlation between ligand–receptor (L–R) interaction scores in 880 and control groups. Each dot represents an L–R pair, with the ligand derived from a gut cell type and the receptor from a brain cell type. Differential interaction levels are calculated as the L–R score in 880 minus the L–R score in control. Pearson correlation coefficient and associated *P*-value are displayed. **f, g**, Number of significantly differential L–R interaction pairs in gut (**f**) and brain (**g**) clusters/cell types. Only statistically significant interactions (*P*-value < 0.05) with both ligand and receptor genes expressed in more than 10% of nuclei are included. Clusters with fewer than 10 differential L–R pairs are omitted from the figure. **h**, Chord diagram showing top-ranked, significantly upregulated (left) and downregulated (right) L–R interaction pairs in 880 compared to control. Edge thickness is proportional to the interaction score, and edge color corresponds to the cluster ID in the gut or brain. **i**, Expression profiles of receptor genes related to EGFR and FGFR signaling pathways across all brain clusters. Notably high expression is observed in glial cell clusters. Circle size represents the proportion of single nuclei expressing each gene, and color intensity indicates normalized expression level. **j**, Single-nucleus level expression comparison of key ligand genes involved in EGFR and FGFR signaling pathways across all gut nuclei from different dietary groups. **k**, Comparison of L–R interaction levels for a curated list of signaling pathway categories across different dietary groups. Only statistically significant interactions (*P*-value < 0.05) are shown. Significance levels were calculated with right-tailed Fisher’s Exact test (**d**), or two-tailed Mann-Whitney test (**j**).

Each of these four selected clusters displayed distinct gene module scores based on canonical cell-type-enriched markers (Fig. 5b), and the cell proportions within these clusters remained comparable between 880-fed and control groups (Fig. 5c). DEGs analysis followed by KEGG pathway enrichment revealed distinct signaling changes within each cluster (Fig. 5d). In both glial clusters, the neuroactive ligand-receptor interaction pathway was among the most significantly upregulated in 880-fed flies. This upregulation was driven by higher expression of *GABA-B-R1*, *Dop2R*, and *Rdl* compared to control and 868-fed flies (Extended Data Fig. 7b, d). All three genes are involved in inhibitory neurotransmission^59^, implying a potential neuroprotective role by reducing excitotoxicity, which would otherwise lead to neuronal damage during aging^60^. In Gamma KCs, folate biosynthesis was significantly upregulated in the 880 group, supported by increased expression of *phu*, *cin*, and *Sptr* (Extended Data Fig. 7c). This suggests enhanced one-carbon metabolism and methylation capacity, processes critical for maintaining cognitive function during aging^61^. Conversely, in ensheathing glial cells from 880-fed flies, genes involved in unsaturated fatty acid synthesis (*Desat1*) and long-chain fatty acid elongation (*Baldspot* and *Acsl*) were significantly downregulated (Extended Data Fig. 7b). Ensheathing glia, which envelop the neuropil and axon tracts in the *Drosophila* central nervous system, may thus undergo a metabolic shift toward accumulating short-chain saturated lipids, consistent with the lipid profile of the 880 yeast strain. This shift could stabilize neuronal membranes and facilitate lipid transfer, providing additional neuroprotection. Furthermore, glycolysis/gluconeogenesis were significantly downregulated in Gamma KCs and perineurial glial cells, while ribosome activity showed downregulation in Alpha/beta KCs (Fig. 5d and Extended Data Fig. 7c-e).

Extending this analysis across all 22 brain clusters revealed a broader pattern: pathways associated with ribosome activity, glycolysis/gluconeogenesis, amino acid biosynthesis, and oxidative phosphorylation (OXPHOS) were significantly downregulated in over half of the clusters in 880-fed flies (Extended Data Fig. 7f and Supplementary Table 4). In contrast, neuroactive ligand-receptor interaction consistently ranked among the top upregulated pathways. Similar trends were observed in the comparison between 880- and 868-fed groups (Extended Data Fig. 7g, h). This metabolic downscaling paired with signaling upregulation suggests a compensatory adaptation wherein neurons and glia may reduce metabolic demands while enhancing communication pathways to preserve neural function in the face of energy decline. Interestingly, Alpha/beta KCs (cluster B8) were the only subpopulation to show upregulation of OXPHOS (Fig. 5d, Extended Data Fig. 7e and Supplementary Table 4). These cells rely heavily on OXPHOS to meet the high energy demands of long-term memory formation and synaptic plasticity^57^, and this capacity is known to decline with age^62^. Thus, our data suggest that dietary lipids from the 880 strain orchestrate a dual metabolic strategy in the aging fly brain: a global downregulation of metabolic activity across most neurons and glia to conserve energy and boost signaling-centric interactions, coupled with a targeted enhancement of mitochondrial function specifically in memory-critical Alpha/beta KCs. This selective metabolic reprogramming likely contributes to cognitive resilience during aging.

While these neural adaptations highlight brain cell-intrinsic metabolic remodeling in response to 880-derived lipids, we next investigated whether systemic signals from the gut also contribute to brain health, given the growing recognition of the gut-brain axis in modulating neuronal function, metabolism, and cognition during aging^39,40^. To explore this, we performed a comprehensive ligand–receptor (L–R) interaction analysis using FlyPhoneDB^63^, designating gut cells as the signal-sending source and brain cells as the signal-receiving targets. In both 880-fed and control flies, we observed largely similar L–R interaction patterns, with comparable interaction scores across different gut and brain clusters (Extended Data Fig. 8a). Consistently, the L–R scores between the two groups were highly correlated (Fig. 5e), suggesting that the majority of gut-brain communication pathways are maintained. However, a number of outlier interactions deviated from the trend line in 880-fed flies, reflecting selective alterations in gut-brain signaling. We grouped these significantly differential L–R pairs by their corresponding gut cell types and found that the number of differential pairs was not proportional to gut cluster size (Fig. 5f). Specifically, as the largest subpopulation, digestive enterocytes in cluster G0 showed relatively few differential interactions, whereas absorptive enterocytes (G1), high-metabolic enterocytes (G2), metal transport enterocytes (G4), and crop cells (G5) contributed disproportionately more differential signals. This finding suggests that 880-derived lipids reshape gut-brain communication primarily through cell types involved in nutrient sensing, metabolism, and stress responses, rather than basic digestive activity. On the brain side, larger clusters such as B0–B5 exhibited higher numbers of differential L–R interactions (Fig. 5g).

Examining the top-ranked, significantly upregulated interactions in 880-fed flies, we found that most signals originating from various gut cell types were directed toward glial cell clusters, particularly B3, B5, and B13, primarily via EGFR and FGFR signaling pathways (Fig. 5h). These involved L–R pairs such as *vn*–*Egfr*, *pyr*–*htl*, and *bnl*–*htl*. We further confirmed that the corresponding receptor genes were exclusively expressed in glial clusters (Fig. 5i). Among the receiving clusters, perineurial glia in cluster B5 represents the dominant target. In *Drosophila* brain, these glial cells form the outermost layer of the blood–brain barrier (BBB) and are instrumental in nutrient uptake and transfer from the hemolymph into the central nervous system^64^. Additionally, EGFR and FGFR signaling are known to regulate glial development, differentiation, and function^65–67^. Thus, the upregulated activation of EGFR/FGFR pathways in perineurial glia following 880 lipid supplementation may strengthen BBB integrity and enhance nutrient regulation, ultimately supporting neural homeostasis and brain health. Moreover, we found significantly increased expression of EGFR/FGFR ligands in 880-fed gut cells, while receptor levels remained comparable across brain groups, suggesting that this diet-responsive modulation is primarily driven by changes in the gut (Fig. 5j and Extended Data Fig. 8b). In contrast, we observed significant downregulation of L–R interactions originating from enteroendocrine cells in cluster G10 via the AstA signaling pathway, targeting multiple neural clusters in the brain (Fig. 5h). Since enteroendocrine cells are major sources of gut-derived hormones that influence brain activity^68^, reduced AstA signaling implies that 880 lipids may modulate gut hormone secretion or alter brain receptor sensitivity, potentially rebalancing gut-brain hormonal crosstalk. A parallel analysis comparing 880- and 868-fed flies revealed a similar pattern of upregulated EGFR/FGFR signaling and downregulated AstA signaling (Extended Data Fig. 8c, d).

To systematically evaluate changes across broader signaling landscapes, we analyzed gut-brain L–R communications encompassing 22 major interactive categories (Fig. 5k). The analysis revealed an overall significantly enhanced interactions between gut cells and glial cells in 880-fed flies, particularly in key pathways such as HIPPO signaling, NOTCH signaling, and positive regulation of guanylate cyclase activity. In contrast, gut-neuron interactions showed few significant differences between dietary groups, with the majority of pathways remaining comparable. Altogether, these findings suggest that 880 yeast-derived lipids reshape gut-brain communication networks to promote protective glial responses in the brain, primarily through the EGFR/FGFR signaling pathways.

### FLY-MAP identifies cell type-specific metabolism and gut metabolic coordination in 880-fed flies

Given the widespread metabolic remodeling observed across gut and brain tissues, we next aimed to systematically characterize the underlying metabolic states at single-cell resolution. To this end, we developed a computational pipeline, FLY-MAP (Fly Metabolic Analysis Pipeline), to extract and analyze metabolic profiles from our snRNA-seq datasets (Fig. 6a). Specifically, we constructed a metabolic matrix by extracting genes associated with Drosophila KEGG metabolic pathways, encompassing 11 major categories and 81 pathways (Supplementary Table 5), from the original gene expression matrix. Unsupervised clustering based on this metabolic matrix was then performed separately for gut and brain nuclei, enabling an unbiased identification of metabolic programs across tissues and dietary conditions.

**Fig. 6:**
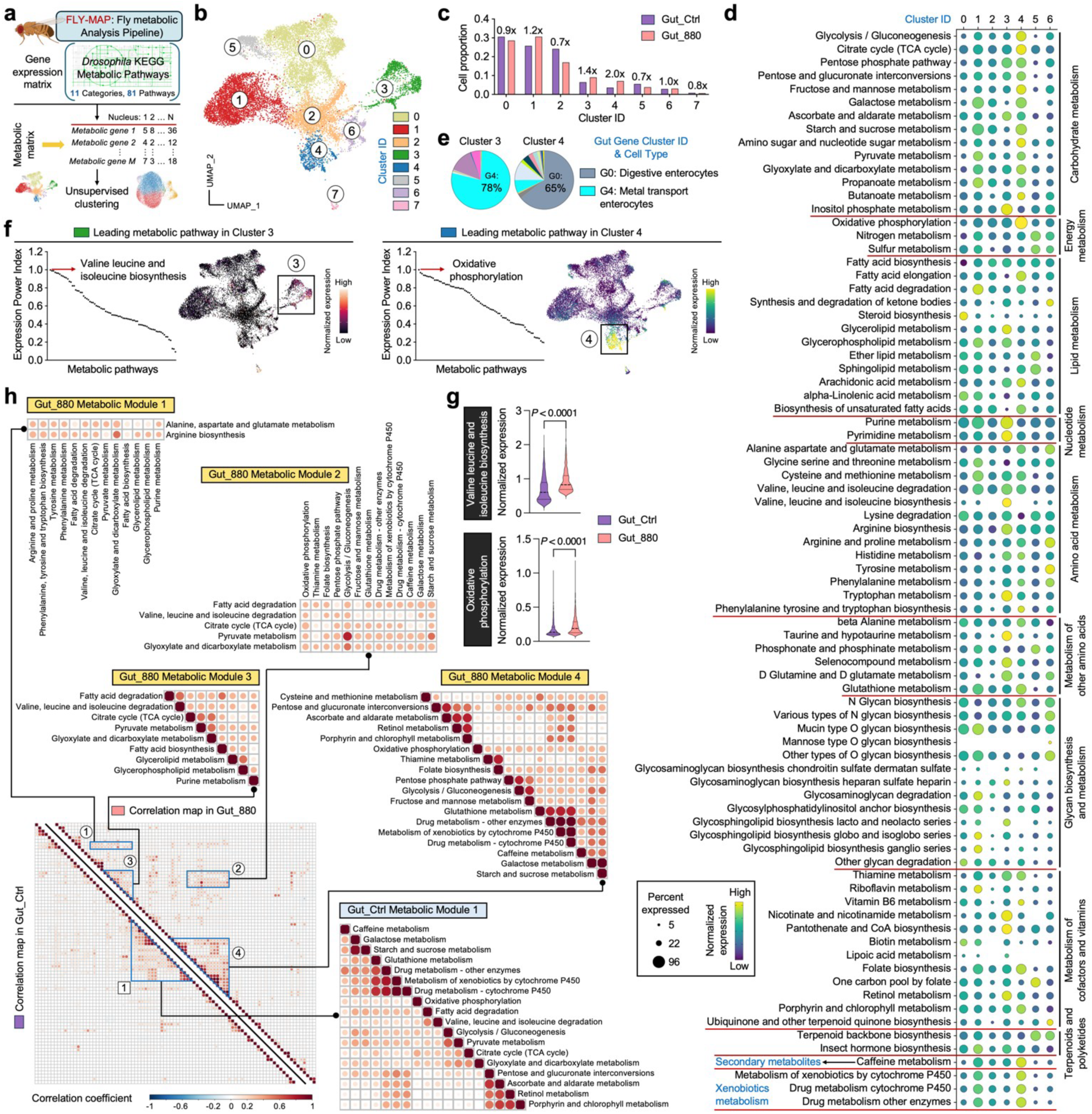
FLY-MAP reveals cell type-specific metabolic states and coordination in 880-fed fly guts. **a**, Schematic of the FLY-MAP (Fly Metabolic Analysis Pipeline). The metabolic matrix was generated by extracting 4 genes associated with *Drosophila* KEGG metabolic pathways from original gene expression matrix, followed by 5 unsupervised clustering performed separately for the gut and brain. Created with BioRender.com. **b**, Unsupervised 6 FLY-MAP clustering of gut nuclei from 880-fed and control flies. **c**, Cell proportions in 880 and control for all 7 identified clusters in panel **b**. Fold changes in cell ratios between 880 and control are indicated for each cluster. **d**, 8 Expression profiles of activity scores of all 81 metabolic pathways across major clusters. Circle size represents the 9 proportion of single nuclei expressing each pathway, and color intensity indicates normalized expression level. **e**, 1 Gut cell type composition of metabolic clusters 3 and 4. Cell type annotations were directly transferred from Fig. 2 3b. **f**, Identification of the leading metabolic pathway in clusters 3 and 4, along with its expression distribution on 3 the UMAP. Pathways are ranked based on the Expression Power Index, defined as the product of expression 4 frequency and expression level. **g**, Single-nucleus level comparison of activity scores for the leading metabolic 5 pathways identified in panel **f** between all nuclei from 880 and control. **h**, Correlation matrices showing expression 6 relationships among all 81 metabolic pathways across all gut nuclei from 880 or control. Pathways are ordered 7 based on hierarchical clustering. Correlation modules with coordinated expression are enlarged for visualization. 8 Color intensity reflects Pearson’s correlation coefficients. Only statistically significant correlations (*P*-value < 0.05) 9 are indicated by dots. Significance levels were calculated with two-tailed Mann-Whitney test (**g**).

In gut nuclei from 880-fed and control flies, FLY-MAP clustering analysis identified 8 distinct metabolic clusters (Fig. 6b). The proportion of cells in each cluster was calculated and compared between the two groups, revealing that cluster 3 had 40% greater occupancy in the 880 group, while the ratio in cluster 4 doubled (Fig. 6c). The stability of other clusters suggests that 880 feeding triggers focused metabolic reprogramming without widespread disruption. In the holistic map of all 81 metabolic pathways categorized by functional class, cluster 3 cells were characterized by elevated nucleotide metabolism, amino acid metabolism, and cofactor and vitamin metabolism, while the higher carbohydrate and energy metabolism in cluster 4 cells suggests they are primed for efficient ATP generation (Fig. 6d). This targeted metabolic remodeling likely supports biosynthesis, stress resilience, redox balance, and energy homeostasis in the 880-fed gut. To resolve cell type identities within each metabolic cluster, FLY-MAP transferred the cell-type annotations from our prior whole-transcriptome clustering (Fig. 3b). Notably, the largest metabolic cluster 0 was dominated by high-metabolic enterocytes, reinforcing the reliability of this independent analysis (Extended Data Fig. 9a). Cluster 3 was primarily composed of metal transport enterocytes (78%), while cluster 4 was largely comprised of digestive enterocytes (Fig. 6e). We next ranked all metabolic pathways in clusters 3 and 4 using the Expression Power Index, defined as the product of gene expression frequency and expression level (Fig. 6f). In cluster 3, the top-ranked pathway was valine, leucine, and isoleucine biosynthesis—the branched-chain amino acid (BCAA) pathway. The gene module score defining this pathway showed notably enriched expression in this cluster, with significantly higher expression observed in 880-fed flies (Fig. 6g). BCAAs not only serve as essential substrates for protein synthesis but also play critical roles in redox regulation and energy metabolism^69^. Their elevated biosynthesis in metal transport enterocytes may support the high metabolic demands of iron handling and contribute to intestinal homeostasis^70^, possibly playing a role in lifespan extension. In cluster 4, OXPHOS was the leading pathway, with significantly increased expression in the 880 group (Fig. 6f, g). Upregulated OXPHOS in digestive enterocytes may fulfill the energy demands required for efficient nutrient absorption and maintenance of epithelial barrier integrity. We also evaluated the metabolic profile of 868-fed flies, revealing an overall dampened metabolic activity compared to the 880 group, particularly in lipid metabolism, nucleotide metabolism, and glycan biosynthesis and metabolism (Extended Data Fig. 9b), potentially explaining their limited ability to extend lifespan.

Motivated by the divergent expression trends across metabolic categories, we next hypothesized that certain pathways may be co-regulated and that dietary lipids may modulate these coordinated interactions. To test this, we incorporated a global correlation analysis into FLY-MAP, comparing pathway relationships in 880-fed and control gut nuclei across all 81 metabolic pathways (Fig. 6h). In control flies, we identified a single large metabolic module that included energy metabolism, redox pathways, xenobiotic metabolism, and other related processes. In gut epithelial cells, glycolysis and fatty acid oxidation generate ATP but also produce reactive oxygen species (ROS). To counteract ROS, cells rely on antioxidant systems, including the glutathione pathway, retinol metabolism, and the pentose phosphate pathway^71^. Thus, the strong correlations observed among energy-generating and redox pathways reflect a fundamental strategy for maintaining cellular homeostasis. In 880-fed flies, we detected the same core module, designated as Metabolic Module 4, supporting its role as a conserved metabolic framework. However, 880 lipids uniquely induced three additional modules (Modules 1–3) defined by significantly positive correlations among glycolysis, BCAA biosynthesis, the TCA cycle, and lipid metabolism. These pathways together orchestrate metabolic flexibility, allowing cells to switch between glucose, amino acids, and fatty acids as energy sources^72^. This flexibility enhances cellular adaptation, stress resistance, and energy balance—processes that are known to decline with aging^73^. Thus, the emergence of these new modules in 880-fed gut may provide a mechanistic basis for their improved metabolic performance and healthspan.

We also applied FLY-MAP to brain tissue, but clustering revealed limited metabolic heterogeneity (Extended Data Fig. 9c). Most brain nuclei were grouped into two dominant clusters with minimal UMAP separation. PCA showed an “elbow point” at ∼10 PCs, indicating that the main metabolic variation was captured in a small number of dimensions (Extended Data Fig. 9d). In contrast, the gut dataset had an elbow at ∼30 PCs, suggesting elevated metabolic complexity and diversity. This difference likely accounts for the reduced cluster resolution in the brain. Further analysis showed similar cell proportions between 880 and control groups across all brain metabolic clusters (Extended Data Fig. 9e). Additionally, metabolic correlation map analysis revealed a similar pattern between the 880 and control groups, with only a single small coordination module identified in both conditions (Extended Data Fig. 9f). All these data point to minimal direct metabolic effects in the brain, suggesting that the metabolic benefits of 880-derived lipids are primarily localized to the gut. Potential improvements in brain function are more likely mediated indirectly through systemic signaling pathways, such as the EGFR and FGFR pathways identified in the previous section. Taken together, our newly developed FLY-MAP reveals cell type-specific metabolic states and coordination patterns, highlighting enhanced metabolic flexibility and regulatory remodeling in gut cells of 880-fed flies, while only limited alterations are observed in brain tissue.

## Discussion

Aging imposes progressive deterioration on tissue structure and metabolic function, particularly within the intestine—a central organ in nutrient absorption, immune regulation, and systemic homeostasis^74^. Despite growing evidence linking gut health to organismal longevity, the potential of dietary lipids to restore intestinal function and extend lifespan remains poorly defined. In this study, we demonstrate that lipids extracted from a genetically engineered long-lived yeast strain confer marked improvements in gut integrity and extend the lifespan of *Drosophila melanogaster*. Through a multimodal approach integrating Raman-based metabolic imaging, single-nucleus transcriptomics, lipidomics, and computational innovation, we uncover a multifaceted mechanism in which these microbial lipids drive epithelial rejuvenation, reprogram metabolic gene networks, and coordinate cross-organ communication via the gut–brain axis.

Our study reveals that engineered yeast-derived lipids extend lifespan by orchestrating a coordinated enhancement of gut epithelial structure, metabolic activity, and cellular homeostasis. Using DO-SRS microscopy, we visualized a striking restoration of LD abundance and membrane lipid incorporation in aged *Drosophila* midguts supplemented with lipids from the long-lived 880 yeast strain. This increase in de novo lipid synthesis was spatially localized to enterocytes, supporting the notion that exogenous lipids can be actively assimilated and remodeled into functional membrane components in aging intestinal tissues. At the molecular level, snRNA-seq revealed a transcriptional reprogramming of gut enterocytes in 880-fed flies, characterized by elevated autophagy, enhanced protein turnover, and reduction of unsaturated fatty acid biosynthesis. These changes occurred most prominently in a metabolically active enterocyte subpopulation, highlighting the selective responsiveness of high-demand cell types to lipid intervention.

A central mechanistic insight emerging from this study is that the unique pro-longevity effects of the 880 yeast strain are underpinned by a distinct lipid remodeling program that shifts the yeast lipidome toward shorter, more saturated fatty acids and membrane-enriching phospholipids. This observation was initially prompted by unsupervised clustering of snRNA-seq data and was subsequently substantiated by Raman spectral analysis and unbiased, mass spectrometry-based lipidomics. Remodeling of the yeast lipidome may be intrinsically linked to the genetic modifications introduced in the engineered strains. Both 880 and 868 strains, which overexpress Hap4, exhibited enhanced mitochondrial function (Fig. 4f) and a significantly increased abundance of defining lipid species associated with mitochondrial membranes (Extended Data Fig. 6d). HAP serves as the principal regulator of heme biogenesis and mitochondrial function in yeast^75^. These findings suggest that Hap4-driven mitochondrial activation and heme metabolism may play a central role in orchestrating the lipidomic reprogramming.

Cross-organ metabolic coordination is increasingly recognized as a fundamental aspect of healthy aging^76^, with the gut–brain axis playing a particularly pivotal role in linking peripheral nutrient status to central nervous system function^39,40^. In this study, we find that strain 880 lipids not only remodel gut metabolism but also induce transcriptional reprogramming in the aging fly brain. L–R interaction analysis highlighted an enhanced gut-to-glia communication, particularly via EGFR and FGFR signaling pathways. However, our FLY-MAP analysis revealed minimal direct metabolic effects within the brain itself, with limited heterogeneity and only subtle changes in metabolic pathway coordination across conditions. These findings suggest that the metabolic benefits of 880-derived lipids are primarily localized to the gut, and that observed improvements in brain function are likely mediated indirectly through gut-initiated signaling cascades. Together, these data support a model in which the aging gut functions not only as a target of intervention but also as an upstream regulator of neural health through lipid-sensitive communication with the brain.

While aging is often associated with widespread metabolic decline, our findings emphasize that the beneficial regulation of 880-derived lipids are not the result of indiscriminate metabolic activation, but rather stem from metabolically precise, cell-type- and context-specific responses. In the brain, snRNA-seq revealed that most neuronal and glial populations exhibited coordinated downregulation of biosynthetic and energy-intensive pathways. Strikingly, this global metabolic dampening was accompanied by a selective enhancement of mitochondrial activity in memory-critical Alpha/beta Kenyon cells. A parallel principle of specificity was observed in the lipidomic dimension: although the 880 and 868 yeast strains shared broad compositional features, only 880-derived lipids extended *Drosophila* lifespan, pointing to the importance of distinct lipid combinations. Functional supplementation with defined lipids demonstrated that a triacylglycerol containing medium-chain saturated fatty acids had modest effects on its own, while phosphatidylcholines alone were ineffective; however, their combination synergistically recapitulated the lifespan extension seen with 880 supplementation. These findings underscore that biological efficacy is not governed by total lipid abundance or generalized metabolic activation, but instead depends on the precise composition, spatial deployment, and contextual interpretation of lipid signals across tissues.

In summary, this study advances a conceptual framework in which engineered microbial metabolism can be harnessed to generate bioactive lipids that exert tissue-specific and age-sensitive effects across a multicellular organism. Unlike traditional dietary restriction or systemic metabolic interventions, which broadly affect many pathways and cell types, our approach demonstrates that precise tuning of lipid composition can achieve metabolic and structural benefits with cell-type selectivity and minimal collateral disruption. The observation that specific yeast-derived lipid species synergistically promote intestinal homeostasis, rewire gut–brain signaling, and ultimately extend fly lifespan underscores the potential of rationally designed lipid formulations as precision nutritional therapeutics. From a translational perspective, this work opens new opportunities to exploit engineered probiotics, functional foods, or lipid-based supplements to modulate aging trajectories. The cross-kingdom nature of this regulatory axis also raises the intriguing possibility that microbial lipid outputs can be tailored to match host tissue vulnerabilities in age-related diseases. Looking forward, integrating systems and synthetic biology of microbial aging^77^ with host-specific metabolic profiling may enable the development of next-generation dietary interventions that align microbial lipid production with personalized aging resilience.

## Methods

### Yeast strains and culture conditions

*Saccharomyces cerevisiae* strains used in this study were derived from the BY4741 background (*MAT*a *his3Δ1 leu2Δ0 met15Δ0 ura3Δ0*). Engineered strains 868, 880, and 898 were generated as previously described^24^. Full strain details are provided in Supplementary Table 6. Frozen glycerol stocks were revived on yeast extract peptone dextrose (YPD) agar plates and incubated at 30 °C for 2 days, after which plates were stored at 4 °C. To maintain strain viability, cultures were re-streaked onto fresh YPD plates every four weeks. For experimental preparation, a single colony was inoculated into 5 mL of synthetic defined (SD) medium (prepared using CSM powder, Sunrise Science, Cat. No. 1001-100, supplemented with 2% glucose) and incubated at 30 °C with shaking at 250 rpm for 24 hours. Cells were then transferred into either 20 mL SD liquid medium or SD agar plates. Cultures were grown until reaching an optical density at 600 nm of 0.6∼0.8.

### Yeast lipid extraction

Total lipid extraction from yeast *S. cerevisiae* was performed using a modified chloroform–methanol protocol. Yeast cultures were harvested by centrifugation at 1,000 × *g* for 5 minutes at room temperature. The supernatant was carefully removed, and the cell pellets were flash-frozen and stored at –80 °C overnight. Frozen samples were then thawed and resuspended in a mixture of 0.5 mL methanol, 0.25 mL chloroform, and 0.5 mL water. The suspension was subjected to ultrasonication in an ice-water bath for 20 minutes. An additional 0.25 mL chloroform was subsequently added, followed by a second round of ultrasonication for 10 minutes. The mixture was centrifuged at 4,000 × *g* for 5 minutes at 4 °C to achieve phase separation. The lower organic phase, containing extracted lipids, was carefully collected and allowed to evaporate overnight in a chemical fume hood. A membrane-like residue, indicative of total lipid content, was observed at the bottom of the tubes after solvent evaporation.

### Fly husbandry

Wild-type *Drosophila melanogaster* (*W^1118^*, stock #5905) were obtained from the Bloomington Drosophila Stock Center and maintained in-house for multiple generations under standard rearing conditions: 25 °C, 60% relative humidity, and a 12-hour light/dark cycle. Flies were raised on cornmeal-based fly food (Nutri-Fly, Genesee Scientific Corporation, Cat. No. 66-113) following established protocols. For yeast lipid supplementation, extracted lipids were mixed with 500 μL of 0.2 M NaOH and incubated at 75 °C for 5 minutes. The resulting mixture was then combined with 2 mL of cornmeal medium, thoroughly mixed, and used as the food source for lipid-fed flies.

### Drosophila lifespan assay

For lifespan assays, 2-day-old adult flies were collected under CO₂ anesthesia and sorted by sex into designated experimental groups, with 20 flies per vial. Flies were maintained on the indicated diets and transferred to fresh food every two days to ensure consistent nutritional conditions. Mortality was recorded daily, and deceased individuals were removed during each transfer. Lifespan data were compiled and used to generate survival curves for statistical analysis.

### Stimulated Raman scattering (SRS) microscopy

SRS imaging was performed using an upright laser-scanning multiphoton microscope (DIY multiphoton, Olympus) equipped with a 25× water immersion objective (XLPLN WMP2, 1.05 NA, Olympus) optimized for near-infrared throughput. The system was integrated with a picoEmerald laser source (Applied Physics & Electronics), providing synchronized picosecond pulsed beams: a tunable pump beam (720–990 nm, 5–6 ps pulse width, 80 MHz repetition rate) and a fixed-wavelength Stokes beam (1032 nm, 6 ps pulse width, 80 MHz repetition rate). Both beams were directed through the sample and collected in transmission using a high numerical aperture oil condenser (1.4 NA).

To isolate the stimulated Raman loss (SRL) signal, the transmitted light was passed through a shortpass filter (950 nm, Thorlabs) to block the Stokes beam while transmitting the pump beam. The filtered pump beam was detected by a silicon photodiode, and the resulting photocurrent was terminated, filtered, and demodulated using a lock-in amplifier operating at 20 MHz. The demodulated signal was processed in real time using the FV-OSR software module (Olympus) integrated with the FV3000 microscope system. Images were acquired at 512 × 512 pixels resolution with a pixel dwell time of 80 μs, corresponding to an imaging speed of approximately 23 seconds per frame. To reduce background interference, reference background images were acquired at an off-resonance wavenumber (1900 cm⁻¹) and subtracted from all SRS images using Fiji (ImageJ).

### Spontaneous Raman spectroscopy

Raman spectra were acquired using a confocal Raman microscope (XploRA PLUS, Horiba) coupled to a spectrometer. A 532 nm diode laser (∼40 mW at the sample) with a line-focus configuration was used as the excitation source. The laser beam was focused onto individual cells using a 100× objective (MPLN100X, Olympus). Laser power was carefully optimized to preserve cell integrity and avoid photodamage. Raman scattering was detected with a thermoelectrically cooled charge-coupled device detector connected to a spectrometer equipped with a 2400 grooves/mm grating. Each Raman spectrum was collected with an integration time of 60 seconds. Instrument calibration was verified using the characteristic silicon peak at 520 cm⁻¹. Background spectra were recorded at the same focal plane for each sample point and subtracted from the raw Raman spectra. All ratio calculations were performed on unprocessed spectral data, prior to any normalization or baseline correction. Spectral data analysis and plotting were carried out using OriginLab software (OriginLab Corporation, Northampton, MA).

### Yeast D_2_O-labeled SRS imaging

To assess metabolic activity via deuterium incorporation, yeast cells were cultured in synthetic defined medium containing 50% D₂O for 12 hours. Following incubation, 5 mL of culture was harvested by centrifugation at 3,000 rpm for 5 minutes at 4 °C. The supernatant was gently removed to avoid disturbing the cell pellet. Cells were fixed in 3.8% paraformaldehyde (PFA) for 30 minutes at room temperature. After fixation, yeast cells were immobilized on poly-L-lysine–coated glass slides for 10 minutes to ensure firm adhesion. Subsequent Raman measurements and SRS imaging were performed on the immobilized cells.

### Fly D_2_O-labeled SRS imaging

To assess age-related changes in metabolic activity, adult flies were administered deuterium-labeled food. Flies at 20 days (middle age) and 40 days (old age) were transferred to fly food containing 20% D₂O and maintained for 5 days. At the end of labeling, five flies from each group (ages 25 and 45 days, respectively) were randomly selected, sacrificed, and their guts were dissected in phosphate-buffered saline (PBS). Dissected gut tissues were fixed in 4% PFA for 15 minutes at room temperature, followed by three washes with PBS in glass wells. The fixed tissues were mounted between a glass microscope slide and coverslip in PBS, and the edges were sealed with nail polish to prevent dehydration during SRS imaging.

### Smurf assay

Gut barrier integrity was assessed using the Smurf assay, as previously described^16^, with minor modifications. Flies with or without yeast lipid supplementation were transferred to vials containing filter paper saturated with a 2.5% (w/v) FD&C Blue No. 1 dye solution (Sigma) in 5% sucrose. Flies were allowed to feed for 6–8 hours at 25 °C. Following incubation, flies were anesthetized on ice and examined under a stereomicroscope. Individuals exhibiting blue dye outside the digestive tract—indicating dye leakage into the hemolymph and body cavity—were scored as “Smurf” flies. The percentage of Smurf flies was calculated for each condition. Assays were conducted using age-matched cohorts, with 20 flies per group and four biological replicates included per experimental condition.

### Fly tissue dissection for single-nucleus RNA sequencing (snRNA-seq)

Dissection and preparation of fly gut and paired brain tissues for snRNA-seq were performed following the protocol established by the Fly Cell Atlas^38^. Briefly, dissections were carried out in ice-cold Schneider’s Drosophila medium. Immediately after dissection, tissues were transferred into 1.5 mL microcentrifuge tubes containing approximately 200 μL of Schneider’s medium, sealed with parafilm, and flash-frozen in liquid nitrogen for >30 seconds. Samples were then stored at –80 °C until nuclei extraction. For each condition, a total of 80 guts and 150 brains were collected to ensure sufficient nuclei yield for sequencing. All dissections were completed within 1 hour to preserve RNA integrity. Nuclei were extracted from frozen tissues through gentle mechanical dissociation. The resulting nuclear suspension was stained with Hoechst 33342 (ThermoFisher, Cat. No. H3570) at a 1:1000 dilution and incubated for 5 minutes. Hoechst-positive nuclei were then isolated using a BD Influx Cell Sorter.

### snRNA-seq library preparation and sequencing

The snRNA-seq libraries were prepared using the Chromium Single Cell 3’ Library & Gel Bead Kit v3.1 (10x Genomics, Cat. No. PN-1000268) according to the manufacturer’s protocol. A total of 30,000 FACS-sorted nuclei suspended in resuspension buffer were loaded onto a Chromium Next GEM Chip G for partitioning. Inside the microfluidic chip, nuclei were co-encapsulated with barcoded gel beads in nanoliter-scale gel bead-in-emulsions (GEMs), enabling cell lysis and reverse transcription within each GEM. Barcoded cDNA was generated through reverse transcription and amplified via 12 cycles of PCR. Amplified cDNA was then subjected to size selection using 0.6× SPRIselect beads (Beckman Coulter, Cat. No. B23317) and quantified using the Agilent TapeStation with D5000 ScreenTape (Agilent, Cat. No. 5067-5588). Subsequent library construction involved cDNA fragmentation, end repair, A-tailing, adapter ligation, and sample index incorporation, followed by 13 additional PCR cycles. Final libraries were assessed for size distribution and quality control before sequencing. Libraries were sequenced on an Illumina NovaSeq X Plus platform, generating 150 bp paired-end reads. Six libraries were pooled per 1000G flow cell with 1% PhiX spike-in control.

### Liquid chromatography–mass spectrometry analysis

Lipid extraction was performed following the Bligh and Dyer method (1959). Briefly, cell pellets were resuspended in 200 μL of water and transferred to glass vials. A volume of 750 μL of chloroform:methanol (1:2, v/v) was added and vortexed thoroughly. Next, 250 μL of chloroform was added, followed by vortexing, and then 250 μL of deionized water was added and vortexed again. Samples were centrifuged at 3,000 rpm for 5 minutes at 4 °C. The lower organic phase was carefully collected into a new glass vial, dried under nitrogen gas, and stored at –20 °C until analysis. Lipid separation was performed on a C30 reverse-phase column (Accucore C30, 2.6 μm, 2.1 mm × 150 mm, Thermo Scientific) maintained at 45 °C. A binary solvent gradient was used with eluent A (acetonitrile:water, 60:40, v/v) and eluent B (isopropanol:acetonitrile, 90:10, v/v), both containing 10 mM ammonium formate and 0.1% formic acid. The chromatographic gradient was programmed as follows: –3 to 0 min: 30% B (isocratic, for column equilibration); 0–2 min: 30% to 43% B; 2–2.1 min: 43% to 55% B; 2.1–12 min: 55% to 65% B; 12–18 min: 65% to 85% B; 18–20 min: 85% to 100% B; 20–25 min: hold at 100% B; 25–25.1 min: 100% to 30% B; 25.1–28 min: 30% B for re-equilibration. The flow rate was maintained at 260 μL/min, and the sample tray was kept at 10 °C. Mass spectrometry was performed on a Q Exactive Orbitrap (Thermo Fisher Scientific) operated in full MS scan mode (resolution 70,000 at m/z 200) followed by data-dependent MS/MS (ddMS², resolution 17,500) in both positive and negative ionization modes. The AGC (automatic gain control) target was set to 1 × 10⁶ for full MS and 1 × 10⁵ for MS/MS. The maximum injection times were 200 ms (MS) and 50 ms (MS/MS). Higher-energy collisional dissociation was performed with stepped collision energies of 25% and 30% in positive mode, and 30 ± 10% in negative mode, using an isolation window of 1.5 Da. Data analysis was conducted using LipidSearch software V4.2.21 (Thermo Fisher Scientific). Only lipid species with molecular identification confidence grade A or B were included in downstream analysis (A: lipid class and all fatty acid chains identified; B: lipid class and partial fatty acid identification). Relative abundances were calculated as the ratio of each lipid species’ peak intensity to the total peak intensity within each yeast strain group.

### Targeted dietary lipid supplementation assay

To assess the functional effects of targeted dietary lipid supplementation, flies were fed cornmeal-based diets mixed with specific short- and medium-chain saturated fatty acids. Four phosphatidylcholine (PC) species, PC(4:0), PC(6:0), PC(10:0), and PC(12:0) (Avanti Polar Lipids, Cat. Nos. 850303, 850305, 850325, 850335) were mixed in equal molar proportions and incorporated into the fly food at a final total concentration of 5 mM. For triglyceride (TG) supplementation, TG(8:0_8:0_8:0) (1,2,3-trioctanoylglycerol; Avanti Polar Lipids, Cat. No. 870111) was added to cornmeal at a final concentration of 5 mM. In the co-feeding condition, equal amounts of the PC mixture and TG(8:0_8:0_8:0) were combined and supplemented into the cornmeal to achieve a total lipid concentration of 5 mM.

### SRS data processing and analysis

SRS spectral data were processed using custom scripts developed in MATLAB (MathWorks), incorporating built-in functions for spectral analysis. Briefly, raw Raman spectral files were imported into MATLAB and interpolated at 1 cm⁻¹ resolution to convert the data into a uniform spectral array. Background signals were subtracted, and baseline correction was applied to reduce signal drift. The resulting spectra were then vector normalized and averaged within each group to minimize noise and enhance the signal-to-noise ratio for biomolecular feature comparison. Individual spectral peaks were assigned to specific molecular vibrations corresponding to distinct chemical bonds or functional groups.

### snRNA-seq data processing and analysis

Raw sequencing reads were aligned to the Drosophila melanogaster reference genome (FlyBase r6.31) using a pre-mRNA GTF annotation established by the Fly Cell Atlas. Alignment, barcode processing, and UMI counting were performed with Cell Ranger V6.1.2 (10x Genomics) to generate a digital gene expression matrix. Quality metrics—including mean reads per nucleus, fraction of valid barcodes, and sequencing saturation—were extracted from the Cell Ranger web summary reports. Subsequent data analysis was conducted using the Seurat V4 pipeline^78^. Low-quality nuclei and potential doublets were filtered based on gene count distributions. Specifically, nuclei with 200–1,600 detected genes were retained for brain tissues, whereas the upper threshold was extended to 3,000 genes for gut tissues due to their inherently higher gene content. Standard Seurat workflow functions were applied in the following order: NormalizeData, FindVariableFeatures, ScaleData, RunPCA, and ElbowPlot to determine the dimensionality of the dataset, followed by FindNeighbors, FindClusters (at a resolution of 0.2 for the gut dataset and 0.1 for the brain dataset), and RunUMAP for visualization. Differentially expressed genes (DEGs) were identified using the FindMarkers function through pairwise comparisons of clusters or yeast strain groups. Module scores for predefined gene sets were calculated using the AddModuleScore function to quantify expression signatures across nuclei.

### KEGG pathway analysis

KEGG pathway analysis of DEGs from each cluster or strain condition was performed using the WebGestalt (https://www.webgestalt.org/) platform^43^. Gene Set Enrichment Analysis mode was applied with KEGG as the functional database, using DEG lists along with their corresponding log₂(fold change) values as input. To identify key regulators or “driver genes” within specific signaling pathways, the Network Topology-based Analysis mode of WebGestalt was employed. This analysis used protein–protein interaction data from the BIOGRID database, with the gene set corresponding to a selected KEGG pathway entered as input.

### Lipid Ontology (LION) enrichment analysis

We utilized the web-based ontology enrichment tool LION/web (http://www.lipidontology.com/) to identify lipid-associated terms in lipidomes^55^. The “LION-PCA Heatmap” mode was employed for LION term identification. Input lipid expression data were submitted and matched against the LION database, with only successfully matched identifiers included in the analysis. LION/web then selected the most variable LION terms by assessing their enrichment across a defined number of principal components. In our dataset, three PCs explaining ∼80% of the cumulative variance were used, and five clusters were determined via hierarchical clustering. Only statistically significant terms (t-test *P*-value < 0.05) were retained for visualization.

### Ligand–receptor (L–R) interaction analysis

Cell–cell connectivity patterns were analyzed using the R package Connectome V1.0.0^79^ in “custom mapping” mode, based on ligand and receptor expression derived from our snRNA-seq datasets. A high-confidence list of L–R pairs covering major *Drosophila* signaling pathways was curated from FlyPhoneDB^63^. The normalized Seurat object was used as input, with cell cluster identities defining the nodes of the interaction networks. The analysis generated an edge list representing connections between node pairs mediated by specific L–R interactions. For global correlation analysis between different groups, differential interaction levels were calculated as the L–R score in group A minus that in group B. A positive value indicates upregulation in group A, while a negative value indicates downregulation. For visualization, top-ranked L–R pairs were selected from statistically significant interactions (*P*-value < 0.05), where both ligand and receptor genes were expressed in more than 10% of nuclei. The “sources.include” and “targets.include” parameters were used to define the source gut cluster emitting ligand signals and the target brain cluster expressing the corresponding receptor genes. Edge thickness in the network visualizations corresponds to correlation weights, with thicker edges indicating stronger interactions.

### Fly metabolic analysis pipeline (FLY-MAP)

The FLY-MAP metabolic matrix was generated by extracting genes associated with *Drosophila* KEGG metabolic pathways from the original whole-transcriptome gene expression matrix. Briefly, gene matrix transposed format data for *Drosophila melanogaster* KEGG pathways were downloaded. FlyBase gene IDs were then converted to gene names using the biomaRt package^80^, and only metabolic pathways were retained for downstream analysis. Seurat standard clustering was performed using the metabolic gene set as variable features for dimensionality reduction. Pathway activity scores were calculated using the AddModuleScore function. Global correlation analysis was conducted based on pathway module scores across all nuclei from each condition. Pearson correlation coefficients were calculated, and pathways were hierarchically clustered and ordered.

### Statistical analysis

Statistical analyses were performed using Prism V10 (GraphPad). Survival data were analyzed using the Log-rank Mantel-Cox test. For comparisons involving three or more groups, one-way ANOVA with Tukey’s multiple comparisons test was used. For two-group comparisons involving non-parametric data, a two-tailed Mann-Whitney test was applied. Unless otherwise specified, scatter plots display mean ± s.e.m. For single-nucleus level comparisons—typically visualized as violin plots—the normalized expression values of the top 10% of single nuclei from each yeast lipid group were included in the analysis.

## Data availability

Raw and processed sequencing data for this study can be accessed in the NCBI Gene Expression Omnibus (GEO) database under the accession number GSE296279. The following secure token has been created to allow review of record GSE296279 while it remains in private status: udyfakmeffstxar

## Code availability

All original code used for analyzing the snRNA-seq data has been deposited in our GitHub repository (https://github.com/Zhiliang-Bai/Drosophila-longevity). Additional information required to reanalyze the data reported in this paper is available from the lead contact upon reasonable request.

## Acknowledgements

We would like to acknowledge the invaluable contribution from Fine Cody at UCSD Human Embryonic Stem Cell Core Facility to help us with cell sorting. Computational data analysis was conducted with the Yale High Performance Computing clusters. We acknowledge the support from UCSD Startup funds, the Hellman Fellow Award and Sloan Research Fellow Award (to L.S.), as well as the support from U.S. National Institutes of Health including grants R01AG086548, R01GM149976, U01AI167892, 5R01NS111039, R21NS125395 (all to L.S.), U54AG076043, U54AG079759 (all to R.F.), R01AG056440, R01AG068112, R01AG086348, R01GM144595, R01AG093633 (to N.H.).

## Author information

These authors contributed equally: Yajuan Li, Zhiliang Bai.

## Contributions

Conceptualization: L.S., R.F., and N.H.; methodology: Y.L., Z.B., L.S., R.F., and N.H.; investigation: Y.L., Z.B., Y.L., J. R., and Y.L.; formal analysis: Y.L., Z.B., F.G., J.V., H. J., Z.L., and S.S.; resources: Y.L., S.Q., A.W.; funding acquisition: L.S., R.F., N.H., and D.S.-K.; writing–original draft, Z.B. and Yajuan Li; writing–review and editing, L.S., R.F., and N.H. with input from all authors.

## Ethics declarations

### Competing interests

L.S., N.H., and Yajuan Li are inventors of a patent application related to this work. R.F. is scientific founder and adviser for IsoPlexis, Singleron Biotechnologies, and AtlasXomics. The interests of R.F. were reviewed and managed by Yale University Provost’s Office in accordance with the University’s conflict of interest policies.

## Figures and Captions

**Extended Data Fig. 1:**
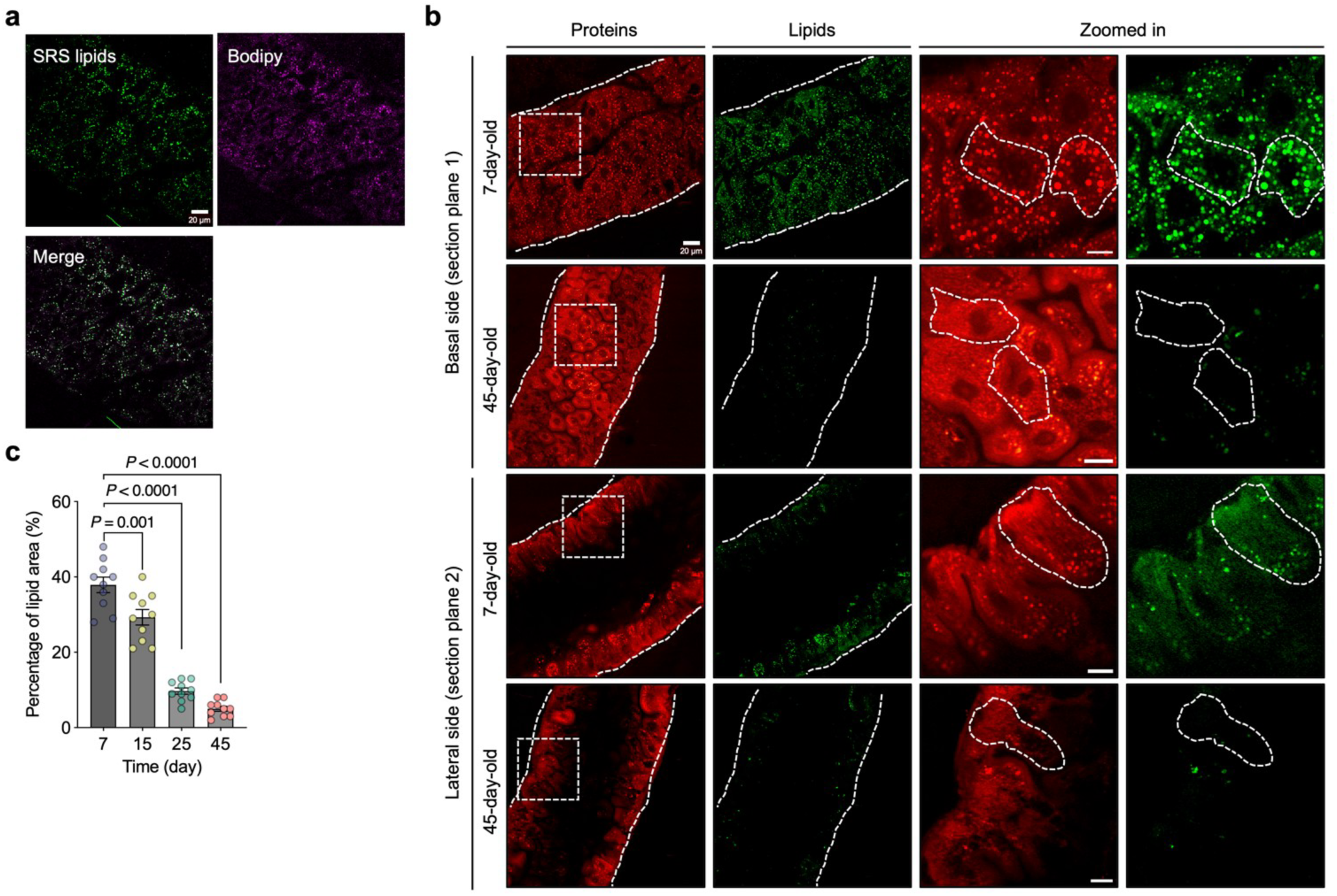
Age-dependent gut lipid alterations visualized by SRS imaging. **a**, Co-registration of BODIPY fluorescence staining with SRS lipid droplet signals in the fly gut, confirming lipid4 specific signal detection. **b**, SRS imaging of gut metabolic profiles in 7-day-old and 45-day-old flies, with separate 5 views of basal and apical-lateral sections. **c**, Quantification of total lipid area in fly guts across different ages. Scatter 6 plot shows mean ± s.e.m. from *n* = 10 flies. Significance levels were calculated with one-way ANOVA with Tukey’s 7 multiple comparisons test.

**Extended Data Fig. 2:**
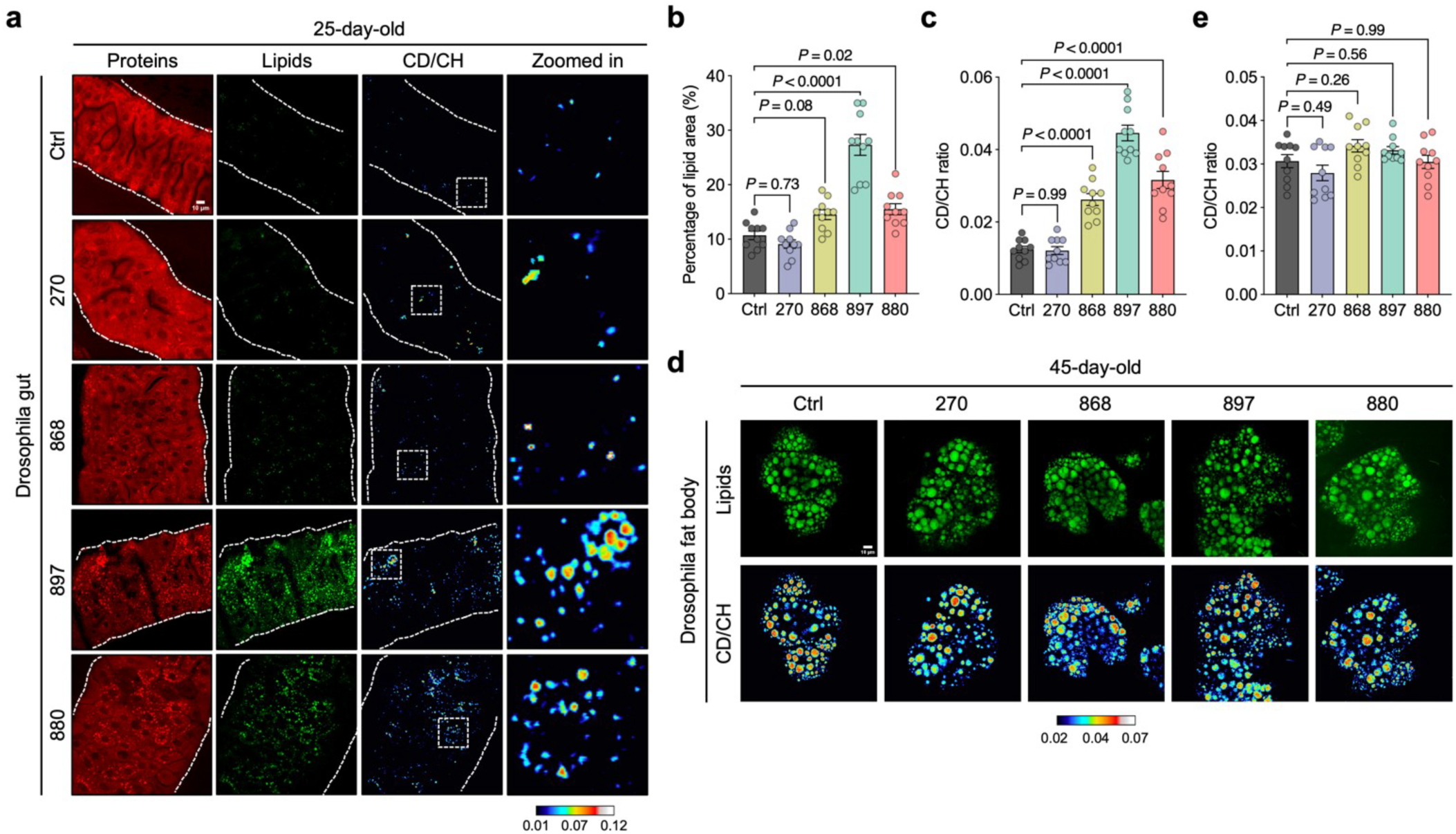
DO-SRS imaging detects lipid metabolism changes in flies fed with different lipid 3 extracts. **a**, DO-SRS imaging of metabolic profiles in the guts of 25-day-old flies fed with different lipid extracts. **b, c**, 5 Quantification of total lipid area (**b**) and CD/CH (2,140/2,850) ratio (**c**) from panel **a** and comparison across yeast 6 strain groups. **d**, DO-SRS imaging of the fat body lipid synthesis in the guts of 45-day-old flies fed with different 7 lipid extracts. **e**, Quantification of CD/CH (2,140/2,850) ratio in panel **d**, compared across yeast strain groups. In **a** and **d**, CD/CH (2,140/2,850) indicates newly synthesized lipids. Scatter plot shows mean ± s.e.m. from *n* = 10 flies (**b**, **c**, and **e**). Significance levels were calculated with one-way ANOVA with Tukey’s multiple comparisons test 10 (**b**, **c**, and **e**).

**Extended Data Fig. 3:**
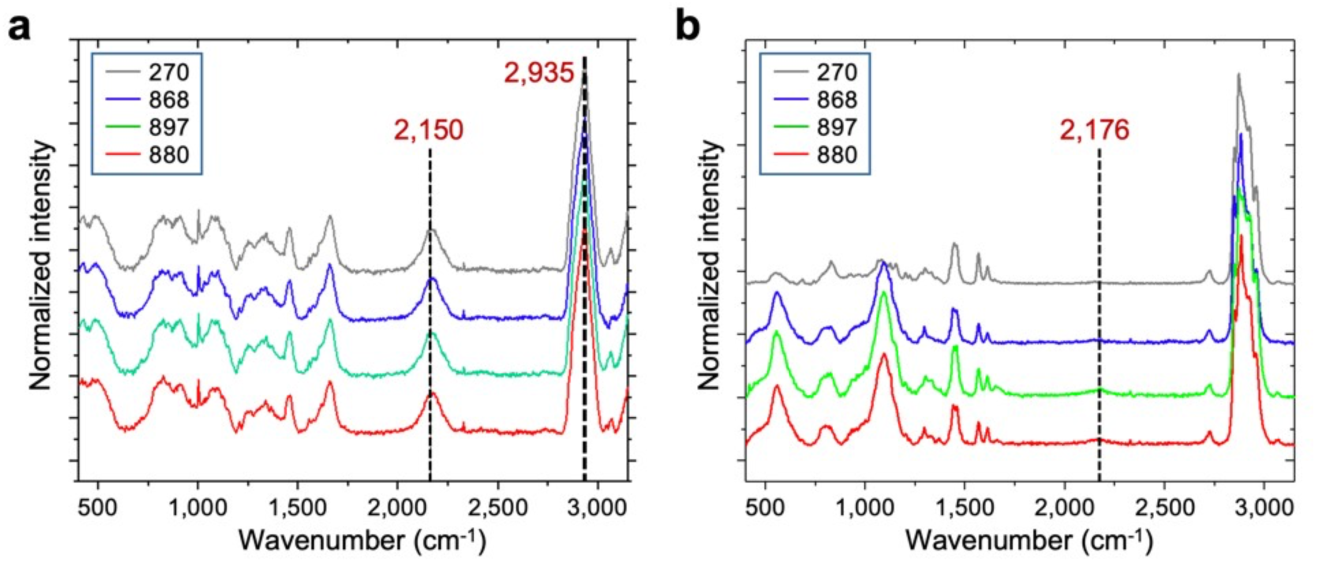
Raman spectra from yeasts and pure lipid extracts. **a**, **b**, Spontaneous Raman spectra of yeast cells (**a**) and pure lipid extracts (**b**) obtained from yeast cultured in D₂O4 labeled media. Each spectrum represents an average from *n* = 10 flies.

**Extended Data Fig. 4:**
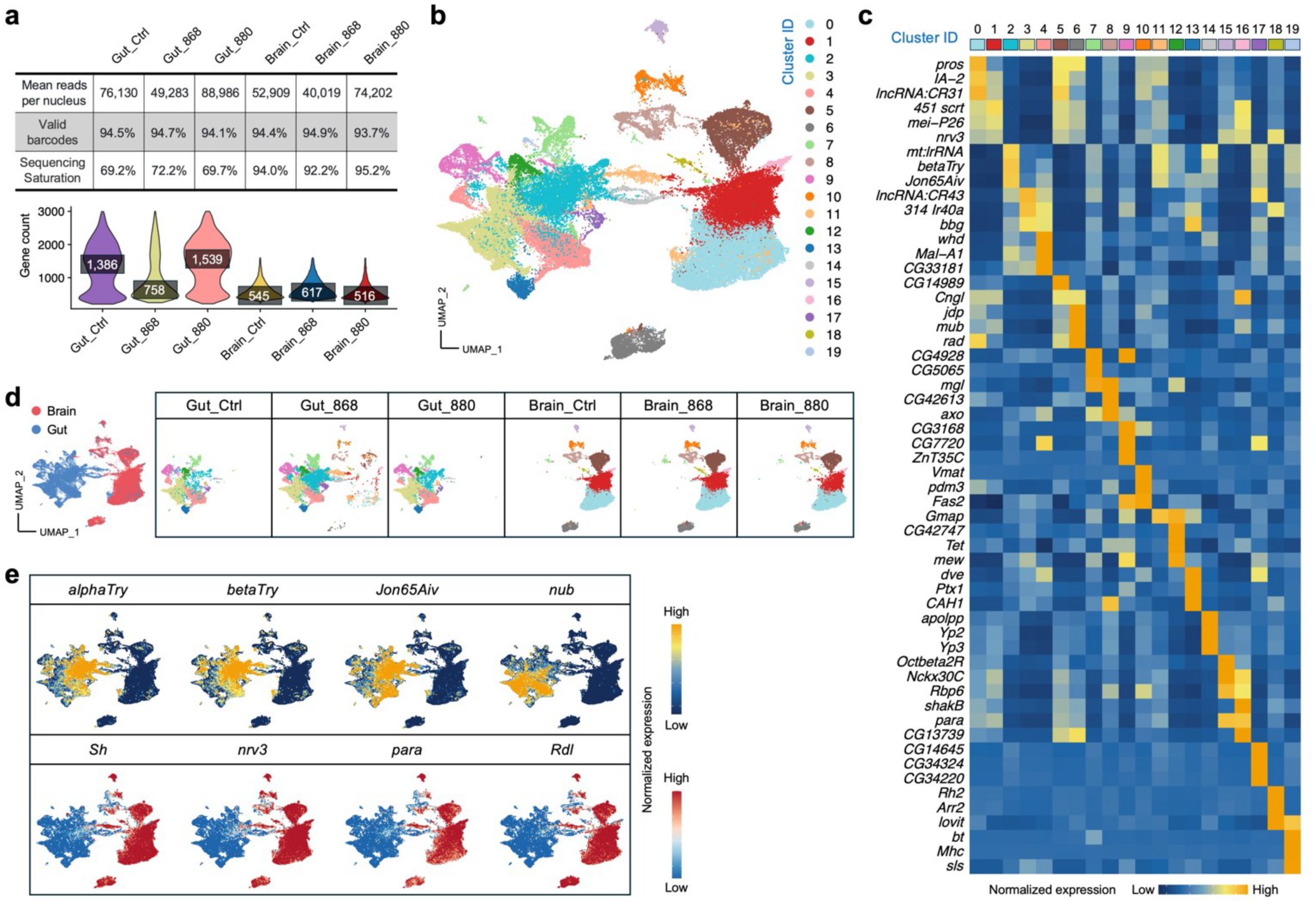
Single-nucleus transcriptomic profiling of fly gut and brain. **a**, Sequencing quality metrics and gene count distributions for each sample. **b**, Integrated UMAP visualization of 4 32,804 gut and 38,277 brain single nuclei from flies fed with different lipid extracts. **c**, Heatmap showing the 5 average expression of the top 3 DEGs across all single nuclei in each cluster identified in panel **b**. **d**, UMAP 6 distribution of all single nuclei grouped by sample source and condition, suggesting minimum batch effect. **e**, 7 Expression distribution of gut- and brain-specific marker genes on the UMAP.

**Extended Data Fig. 5:**
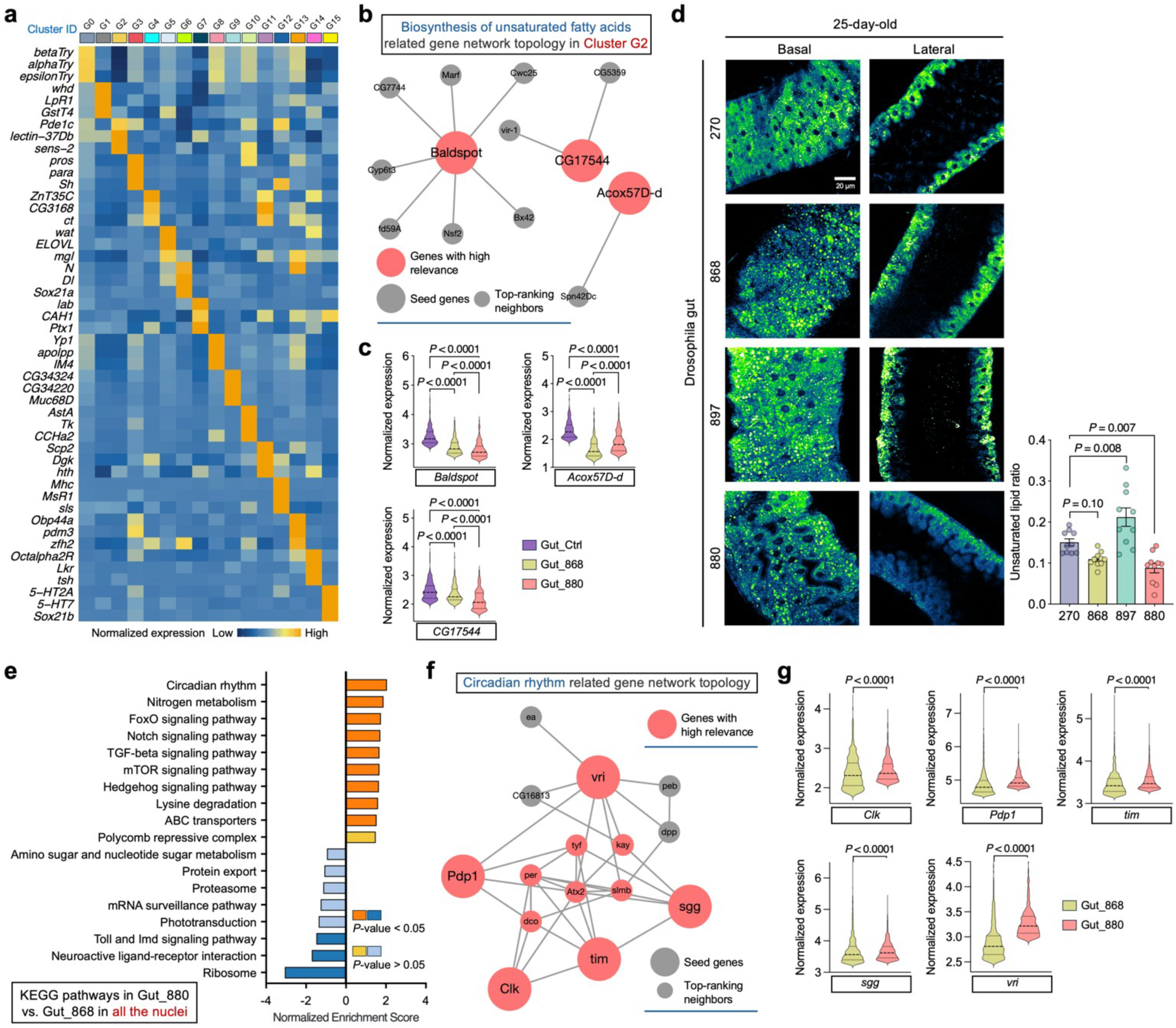
snRNA-seq, SRS imaging, and signaling pathway analysis of fly guts. **a**, Heatmap showing the average expression of the top 3 DEGs across all single nuclei in each cluster identified in Fig. 3b. **b**, Network topology analysis of unsaturated fatty acids synthesis related genes in cluster G2. **c**, Singlenucleus level expression of key genes identified in panel **b** and comparison across dietary groups. **d**, Ratiometric images of the 3,012/2,850 signal on both basal and lateral sides of gut epithelial cells from 25-day-old flies fed with different lipid extracts, accompanied by quantitative comparisons across yeast strain groups. **e**, KEGG signaling pathways enriched among DEGs identified across all single nuclei from 880-fed flies compared to 868-fed flies. **f**, Network topology analysis of circadian rhythm related genes corresponding to panel **e**. **g**, Single-nucleus level expression of key genes identified in panel **f** and comparison between 880- and 868-fed flies. Scatter plot shows mean ± s.e.m. from *n* = 10 flies (**d**). Significance levels were calculated with two-tailed Mann-Whitney test (**c**, **g**), or one-way ANOVA with Tukey’s multiple comparisons test (**d**).

**Extended Data Fig. 6:**
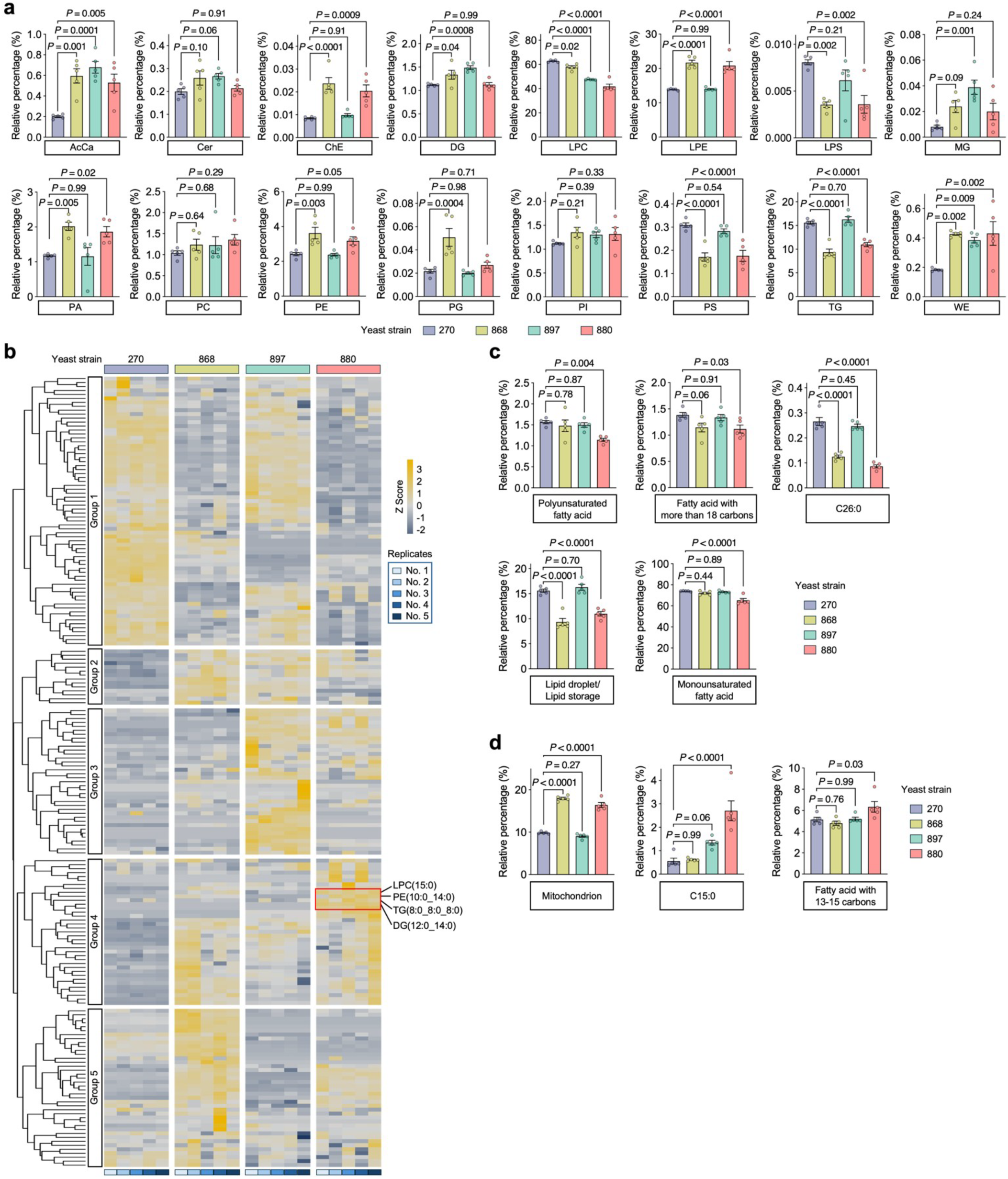
Lipidomics profiling of lipid extracts from the four yeast strains. **a**, Relative abundance comparison of each lipid class across different yeast strains. **b**, Heatmap of scaled lipidomic 4 expression data, with samples on the x-axis and individual lipid species on the y-axis. Lipid species were clustered 1 into five groups using hierarchical clustering. Selected lipids enriched in strain 880 are highlighted. Z-score > 0 2 indicates upregulation; Z-score < 0 indicates downregulation. **c**, **d**, Summed expression levels of key lipid 3 metabolites defining selected lipid pathway terms, compared across yeast strains. Pathways that are upregulated (**c**) 4 or downregulated (**d**) in strain 880 are shown separately. Scatter plot shows mean ± s.e.m. from *n* = 5 replicate 5 measurements (**a, c, and d**). Significance levels were calculated with one-way ANOVA with Tukey’s multiple 6 comparisons test (**a, c, and d**).

**Extended Data Fig. 7:**
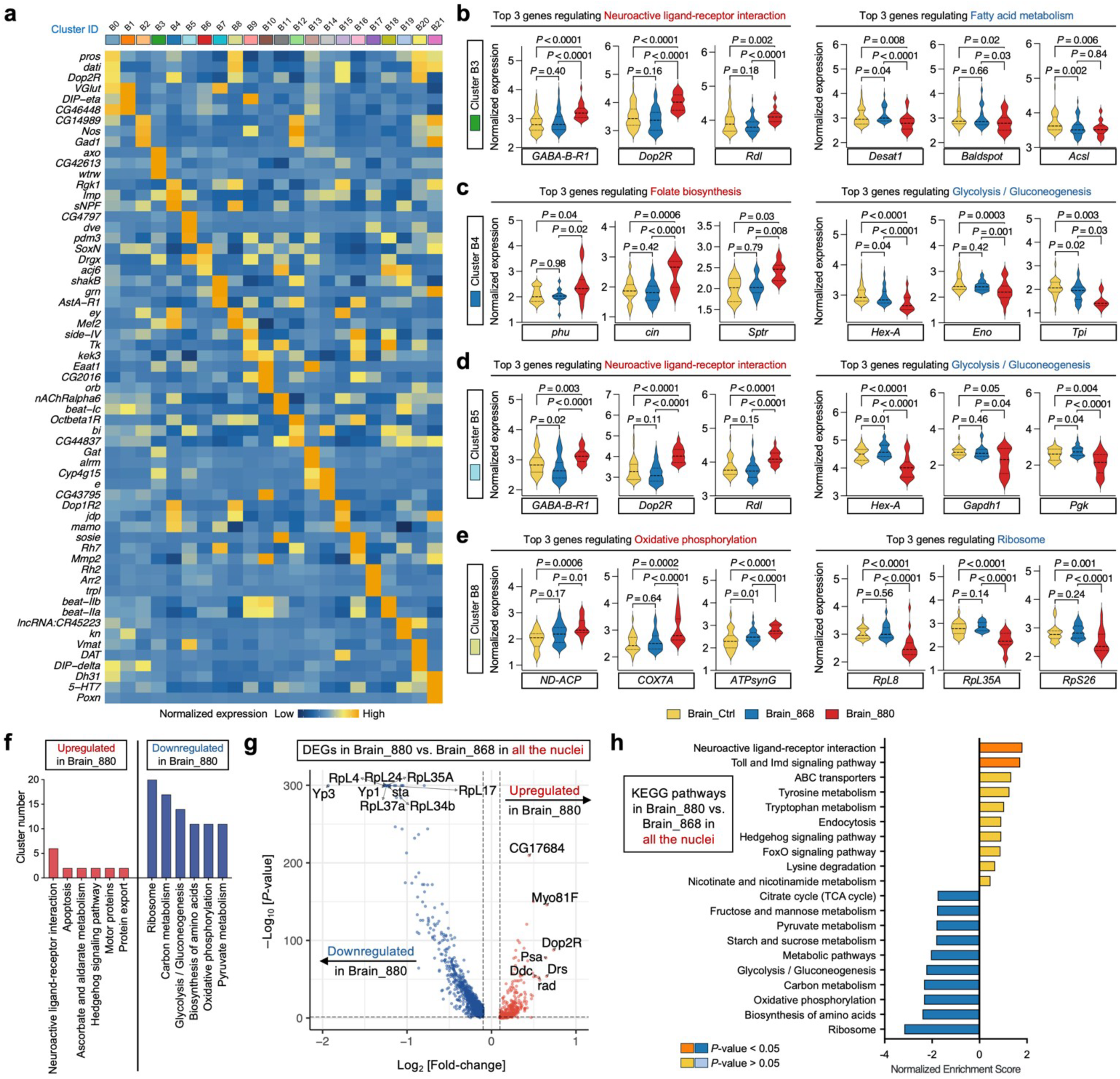
snRNA-seq and signaling pathway analysis of fly brain nuclei. **a**, Heatmap showing the average expression of the top 3 DEGs across all single nuclei in each cluster identified in 4 Fig. 5a. **b**–**e**, Single-nucleus level expression comparison of key genes involved in top-ranked signaling pathways 5 identified in Fig. 5d across cluster B3 (**b**), B4 (**c**), B5 (**d**), and B8 (**e**) nuclei from different dietary groups. **f**, Top6 ranked signaling pathways significantly (*P*-value < 0.05) upregulated or downregulated in 880-fed flies across most 7 identified clusters. **g**, DEGs across all single nuclei in 880-fed flies compared to 868-fed flies. **h**, KEGG signaling 8 pathways regulated by the DEGs identified in panel **g**. Significance levels were calculated with two-tailed Mann9 Whitney test (**b**–**e**).

**Extended Data Fig. 8:**
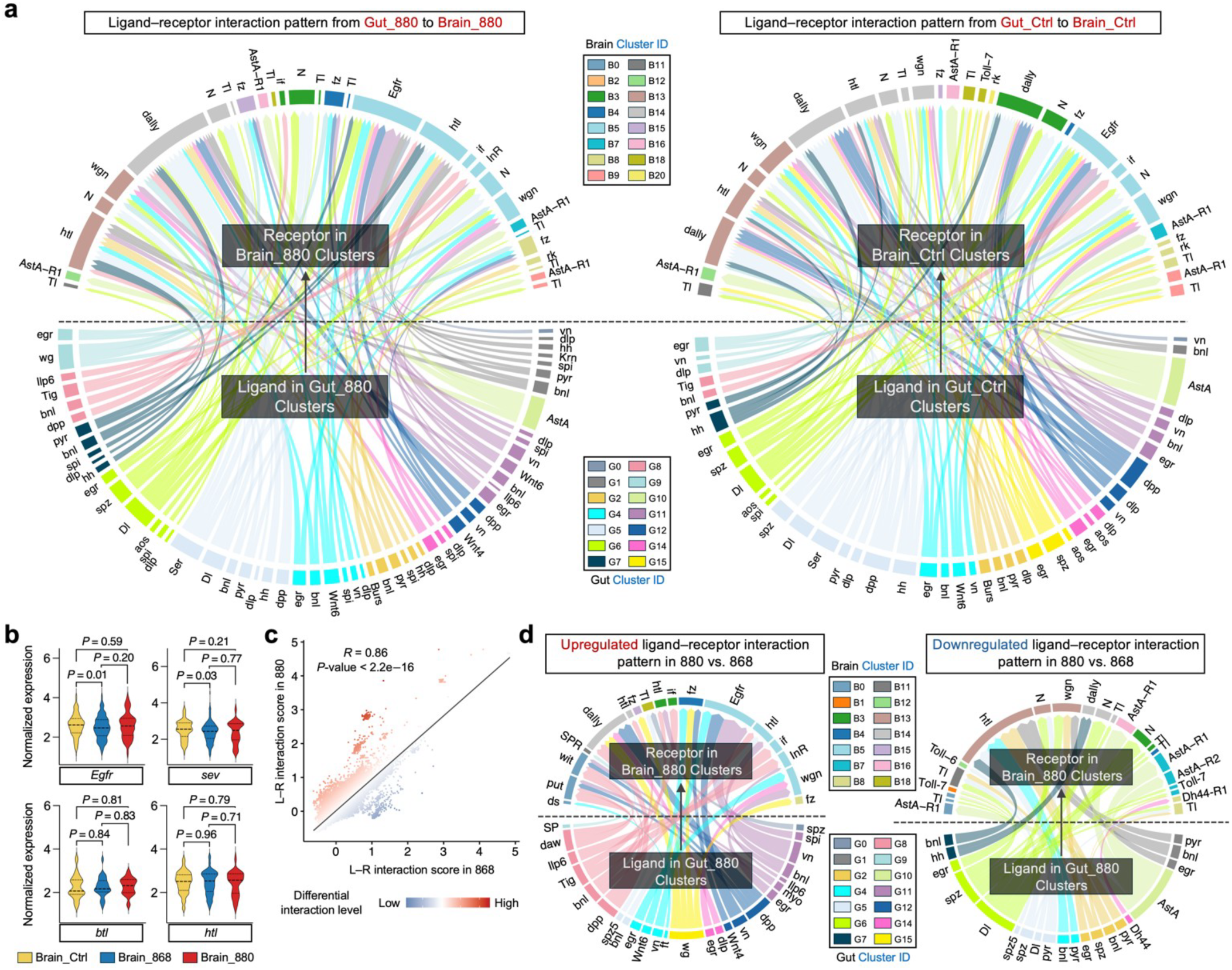
L–R interaction analysis between gut and brain clusters. **a**, Chord diagram showing top-ranked L–R interaction pairs in 880-fed flies (left) and control group (right). Only statistically significant interactions (*P*-value < 0.05) with both ligand and receptor genes expressed in more than 10% of nuclei are included. **b**, Single-nucleus level expression comparison of key receptor genes involved in EGFR and FGFR signaling pathways across all brain nuclei from different dietary groups. **c**, Correlation between L–R interaction scores in 880 and 868 groups. Each dot represents an L–R pair. Differential interaction levels are calculated as the L–R score in 880 minus the L–R score in 868. Pearson correlation coefficient and associated *P*value are displayed. **d**, Chord diagram showing top-ranked, significantly upregulated (left) and downregulated (right) L–R interaction pairs in 880 compared to 868. In **a** and **d**, edge thickness is proportional to the interaction score, and edge color corresponds to the cluster ID in the gut or brain. Significance levels were calculated with twotailed Mann-Whitney test (**b**).

**Extended Data Fig. 9:**
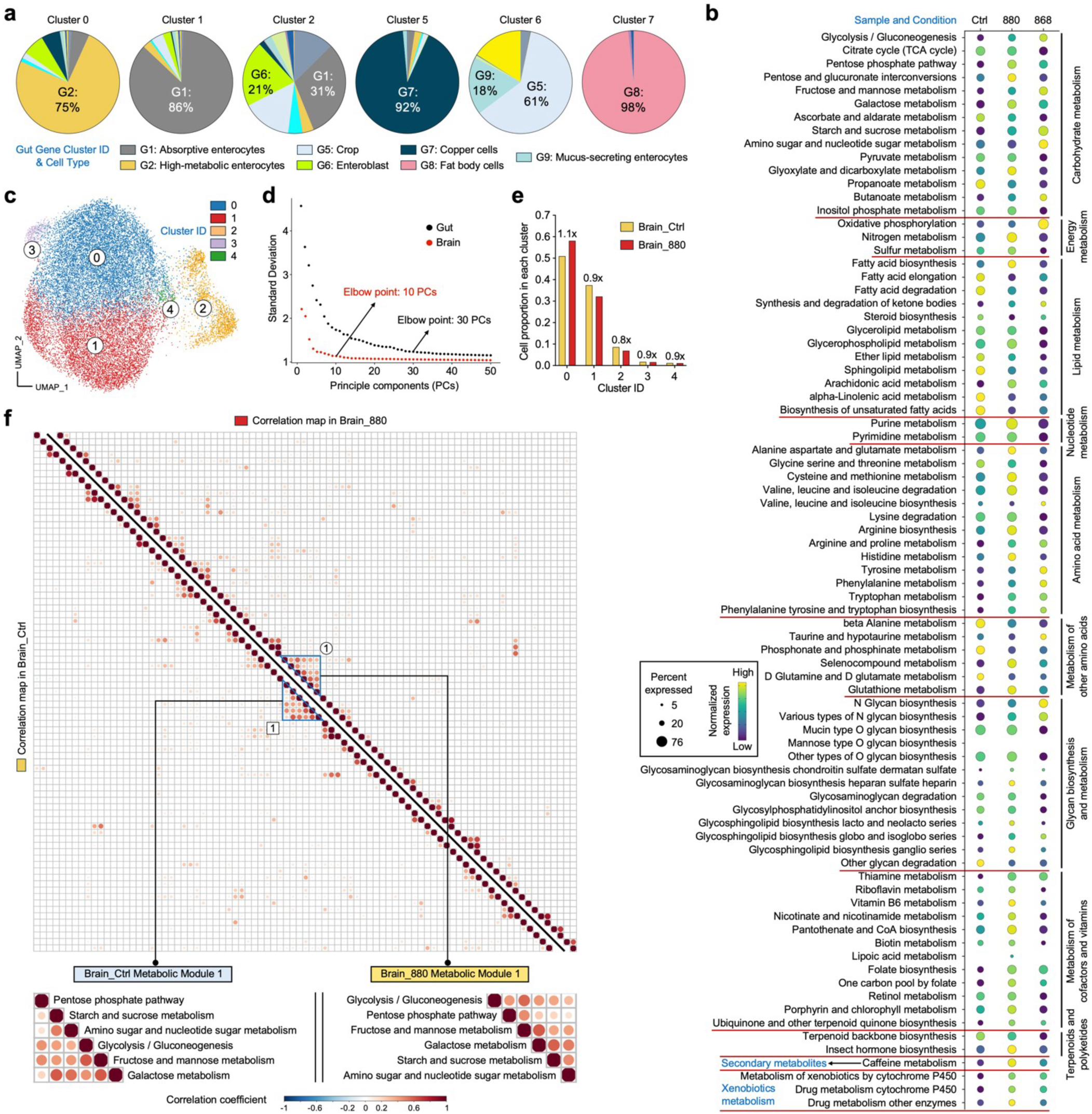
FLY-MAP analysis of gut and brain nuclei. **a**, Gut cell type composition of metabolic clusters 0, 1, 2, 5, 6, and 7. Cell type annotations were directly transferred from Fig. 3b. **b**, Expression profiles of activity scores of all 81 metabolic pathways across all gut nuclei from 5 different dietary groups. Circle size represents the proportion of single nuclei expressing each pathway, and color 6 intensity indicates normalized expression level. **c**, Unsupervised FLY-MAP clustering of brain nuclei from 880-fed 7 and control flies. **d**, Ranking of principal components based on standard deviation in gut and brain datasets. **e**, Cell 8 proportions in 880 and control for all identified clusters in panel **c**. Fold changes in cell ratios between 880 and control are indicated for each cluster. **f**, Correlation matrices showing expression relationships among all 81 10 metabolic pathways across all brain nuclei from 880 or control. Pathways are ordered based on hierarchical clustering. Correlation modules with coordinated expression are enlarged for visualization. Color intensity reflects Pearson’s correlation coefficients. Only statistically significant correlations (*P*-value < 0.05) are indicated by dots.

## Notes

### Competing Interest Statement

The authors have declared no competing interest.

